# Deletion of *Abi3*/*Gngt2* influences age-progressive amyloid β and tau pathologies in distinctive ways

**DOI:** 10.1101/2021.11.08.467701

**Authors:** Kristen R Ibanez, Karen N McFarland, Jennifer Phillips, Mariet Allen, Christian Lessard, Lillian Zobel, Elsa Gonzalez De La Cruz, Shivani Shah, Quan Vo, Xue Wang, Zach Quicksall, Daniel Ryu, Cory Funk, Nilufer Ertekin-Taner, Stefan Prokop, Todd E Golde, Paramita Chakrabarty

## Abstract

The S209F variant of Abelson Interactor Protein 3 (*ABI3*) increases risk for Alzheimer’s disease (AD), but little is known about ABI3 function. RNAscope showed Abi3 is expressed in microglial and non-microglial cells, though its increased expression appears to be driven in plaque-associated microglia. Here, we evaluated *Abi3^-^*^/-^ mice and document that both *Abi3* and its overlapping gene, *Gngt2*, are disrupted in these mice. Expression of *Abi3* and *Gngt2* are tightly correlated, and elevated, in rodent models of AD. RNA-seq of the *Abi3*-*Gngt*2^-/-^ mice revealed robust induction of an AD-associated neurodegenerative signature, including upregulation of Trem2, Plcg2 and Tyrobp. In *APP* mice, loss of *Abi3*-*Gngt2* resulted in a gene dose- and age-dependent reduction in Aβ deposition. Additionally, in *Abi3*-*Gngt2*^-/-^ mice, expression of a pro-aggregant form of human tau exacerbated tauopathy and astrocytosis. Further, the AD-associated S209F mutation alters the extent of ABI3 phosphorylation. These data provide an important experimental framework for understanding the role of *Abi3*-*Gngt2* function in AD. Our studies also demonstrate that manipulation of glial function could have opposing effects on amyloid and tau pathology, highlighting the unpredictability of targeting such pathways in AD.

## Introduction

Alzheimer’s disease (AD) is associated with several canonical pathological hallmarks such as extracellular amyloid β (Aβ) deposits, intracellular neurofibrillary tangles (NFT) of tau protein and robust immune activation (Jack et al., 2018). Activation of innate immunity is an integral part of the AD pathological cascade, though its role is not fully understood due to the diversity and complexity of the immune signaling processes (Dansokho & Heneka, 2018, Golde, 2019, Hansen et al., 2018, Karran & De Strooper, 2016).

Recent genetic and transcriptomic data have highlighted the involvement of immune signaling processes in AD pathogenesis (Golde, 2019, Hansen et al., 2018, Karran & De Strooper, 2016, Shi & Holtzman, 2018). Several genes, whose expression is thought to be enriched in microglial cells, have been implicated as AD risk factors (Efthymiou & Goate, 2017, Walker, 2020). One of these recently identified genes encodes the Abl Interactor family member 3 (ABI3), also known as NESH (new molecule including SH3) (Conway et al., 2018, Sims et al., 2017). The S209F *ABI3* variant (rs616338:p.Ser209Phe) increased the risk of AD (OR=1.43, p=4.5×10^-10^, MAF=0.008) (Conway et al., 2018, Sims et al., 2017) and has been validated in a secondary study (Wightman et al., 2021).

The ABI3 protein contains a Src homology 3 (SH3) domain, a homeobox homology domain and several proline-rich and serine-rich motifs (Miyazaki et al., 2000). Its putative function, largely based on analogy to function of other family members, is to regulate actin polymerization resulting in reduced ectopic metastasis of tumor cells and cell migration (Ichigotani et al., 2002). In the periphery, it is highly expressed in spleen, lymph node and appendix. In the brain, its expression has been reported to be enriched in microglia with ramified or amoeboid morphology (Satoh et al., 2017). Single cell data from mouse brain suggests low level expression in microglia ((Yao et al., 2021); data available from: celltypes.brain-map.org). In the brain, it may also play a role in dendritic spine morphogenesis (Bae et al., 2012a). Though the function of ABI3 in Alzheimer’s pathogenesis or any neurodegenerative disease is unknown, co-expression network analysis suggests a close functional relationship with at least two other notable AD related microglial genes, *TREM2* and *SPI1* (Sims et al., 2017).

Here we have evaluated expression of Abi3 in mouse models of AD pathologies and in bulk RNAseq studies of human brain. We find that Abi3 RNA is not just present in microglia but also in non-microglial cells. However, Abi3 mRNA increases seen in human AD and these mouse models appear to be driven by increases in microglia especially those surrounding amyloid plaques. We have also characterized an *Abi3*^-/-^ mouse line (B6N(Cg)-Abi3tm1.1(KOMP)Vlcg/J), and the effects of loss of *Abi3* locus on amyloid and tau pathologies. Initial characterization using bulk RNAseq of the brain reveals loss of expression of an overlapping gene, *Gngt2*, indicating these mice are better referred to as *Abi3*-*Gngt2*^-/-^. Notably, these mice showed early upregulation of an immune gene expression profile, characterized by increased levels of Trem2, Plcg2 and Tyrobp, that has been previously documented to be induced in rodent models of AD pathologies and AD brains. Haploinsufficiency (+/-) or complete knockout (-/-) of the *Abi3*-*Gngt2* locus reduced Aβ levels in an *APP* mouse model. This beneficial effect was diminished as the mice aged. In *Abi3*-*Gngt2*^-/-^ mice, overexpression of a pro-aggregant tau using neonatal delivery of adeno-associated viruses (AAV) resulted in increased tau pathology and astrocytosis. We further find that the S209F ABI3 variant alters phosphorylation of ABI3, providing initial insight into how this mutant may alter ABI3 function. These data further highlight the complexity of immunoproteostasis with respect to AD relevant pathologies (Golde, 2019), and reinforce the notion that factors that alter immune signaling often have opposing effects on amyloid and tau.

## Results

### ABI3 is expressed in microglia and neurons in mice and humans

Recent reports have confirmed *ABI3* as an AD risk gene (Conway et al., 2018, Sims et al., 2017, Wightman et al., 2021). *ABI3* has been reported to be a microglia-specific gene using single cell RNAseq (Yao et al., 2021, Zhang et al., 2014). Bulk RNAseq data reveals increased levels of ABI3 RNA relative to control cohorts in both the temporal cortex and cerebellum of human patients and in mouse models of amyloid and tau pathologies (Sims et al., 2017). In AD brains, ABI3 expression is significantly upregulated (p= 4.47E-03) (Sims et al., 2017). We further confirmed this in aging cohorts of *APP* TgCRND8 (TG) mice relative to nontransgenic (NonTG) age-matched mice (Fig. 1a**; Table EV1**). It is notable that increasing plaque burden, and not necessarily age, seems to be associated with increased Abi3 expression (Fig. 1a). In *MAPT* transgenic rTg4510 mice, the RNA levels of Abi3 increased in 4.5 month old tau expressing mice but reduced to levels comparable to NonTG littermates at 6 months (Fig. 1b**; Table EV2**).

**Fig. 1.**
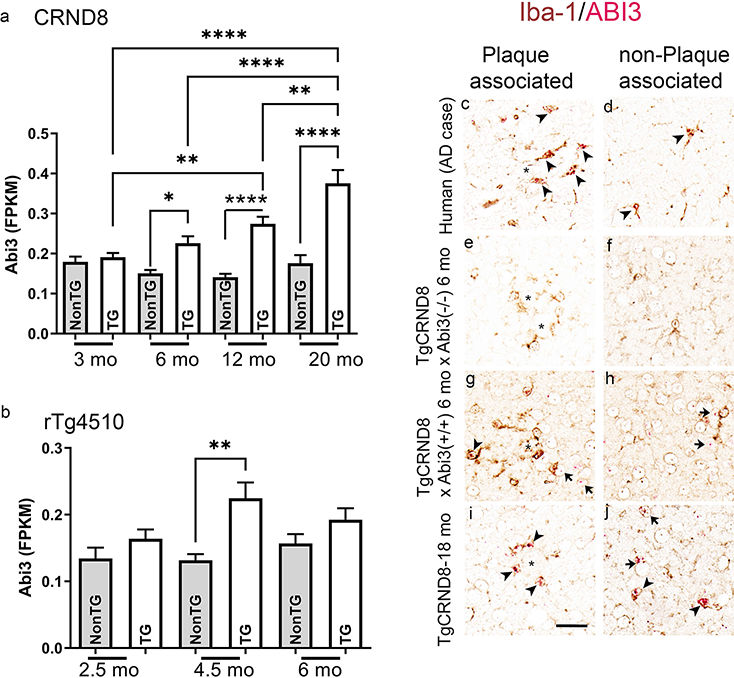
ABI3 RNA is expressed in microglia and neurons in mice and humans. a-b. Abi3 RNA levels (FKPM) plotted across different ages in *APP* CRND8 mice (a) and *MAPT* rTg4510 mice (b). Data obtained from AMP-AD Synapse data from UF-Mayo group. n=9-14 (a) and n=6 (b). 1 way Anova, ****p<0.0001; **p<0.01; *p<0.05. c-j. In situ hybridization was done to detect ABI3 (Fast Red; red color) RNA on human and mouse paraffin embedded brain sections immunostained with anti Iba-1 antibody (brown color). Asterisks mark Aβ deposits; arrowheads indicate Iba-1 (microglia) associated in situ signal and arrows indicate in situ signal in non-microglia cells. n=3 (human AD cases, 6 month old TG-*Abi3-Gngt2^-/-^* mice and 6 month old TG*-Abi3-Gngt2^+/+^* mice) and n=1 (20 month old TgCRND8 mice). Representative of two independent experimental replicates. Additional representative images are available in Fig. EV1. NonTG, nontransgenic; TG, transgenic.

To further evaluate Abi3 expression, we used RNAscope to localize Abi3 expression in mouse and human tissues (Fig. 1c-j**, Fig. EV1**). In AD patients, we observed Abi3 transcript in Iba1-immunopositive microglia abutting Aβ plaques (Fig. 1c; **Fig. EV1a-c**). Interestingly, we also observed Abi3 transcripts in Iba-1 nonreactive cells in the grey matter, which are presumed to be neurons based on size and location (**Fig. EV1d-f**). Specificity for the RNA in situ hybridization signal was confirmed by using brain tissue from *APP* mice completely lacking the *Abi3* locus (Fig. 1e-f; **Fig. EV1g-l**). In 6 month old *APP* TgCRND8 mice, we observed Abi3+ microglia around Aβ deposits in the cortex (Fig. 1g; **Fig. EV1m-n**) and in the white matter tracts (**Fig. EV1o**). Similar to human cases, we also noted Abi3+ cells in the neuronal layers in the cortex and hippocampus (Fig. 1h; **Fig. EV1p-r**). Microglia in close proximity of Aβ plaques showed increased levels of Abi3 RNA in these TG mice (6 month old) (**Fig. EV1m-o**). We observed increased Abi3 positivity in Aβ plaque-associated microglia in 18 month old CRND8 mice (Fig. 1i; **Fig. EV1s-u**). These older mice also showed Abi3 RNA in forebrain neurons (Fig. 1j**; Fig EV1u, x**).

### *ABI3* and *GNGT2* genes are co-regulated in *APP* mouse models and in AD

To elucidate how Abi3-specific immune signaling contributes to the neurodegenerative cascade process, we obtained the *Abi3*^-/-^ mice from Jax Labs (Stock #028180). While using bulk RNAseq to characterize these mice, we serendipitously discovered that the levels of another gene, *Gngt2*, was dramatically reduced to the same extent as *Abi3* (**Fig. EV2a**). We surveyed the gene maps and found that mouse *Abi3* locus overlaps with two other genes on chromosome 11 – microglia-specific G protein gamma transducing activity polypeptide 2 (*Gngt2*) and *Phospho-1* (**Fig. EV2b**). We also confirmed that in humans, the arrangement of the *ABI3*, *GNGT2* and *PHOSPHO-1* genes is conserved, albeit on chromosome 17. Using data from UCSC genome browser, we discovered that the Velocigene targeted deletion in the *Abi3* locus knocked out both *Abi3* and *Gngt2* (**Fig. EV2b**). In subsequent sections, we will refer to these mice as *Abi3-Gngt2*^-/-^. Notably, we did not find any changes in Phospho-1 levels in these mice.

Given that *Abi3* and *Gngt2* genes are overlapping, we hypothesized that their expression could be correlated. Using RNAseq data from the AMP-AD consortium, we investigated the concordance between the expression patterns of *Abi3* and *Gngt2*. We found that *ABI3* and *GNGT2* genes were co-regulated in three distinct AD patient cohorts: temporal cortex samples of Mayo Clinic AD cohort (Mayo TCX: ρ = 0.644, p = 2.2e-16) (Fig. 2a), cerebellar samples of Mayo Clinic AD cohort (Mayo CER: ρ = 0.556, p = 2.2e-16) (Fig. 2b) and Religious Orders Study and Rush Memory and Aging Project AD cohort (ROSMAP; ρ = 0.328, p = 2.2e-16) (Fig. 2c). We also confirmed that Abi3 and Gngt2 genes are co-regulated in the *APP* transgenic TgCRND8 mice (Fig. 2d, ρ = 0.625, p = 7.31-06) and in *MAPT* transgenic rTg4510 mice (Fig. 2e, ρ = 0.554, p = 0.018). The NonTG littermates of these mice showed no correlation between *Abi3* and *Gngt2* expression. In addition, we investigated 96 additional mouse transcriptomic datasets (Al-Ouran et al., 2019, Wan et al., 2020) and in 26 cohorts, we found that *Abi3* and *Gngt2* were both differentially regulated. Among these 26 of these cohorts, *Abi3* and *Gngt2* expression changes were concordant in 24 studies (**Table EV3**), showing that these genes are consistently co-regulated across different mouse models and experimental cohorts. Collectively, these analyses show that expression of *ABI3* and *GNGT2* genes are in a tight co-expression network in AD and AD mouse models.

**Fig. 2.**
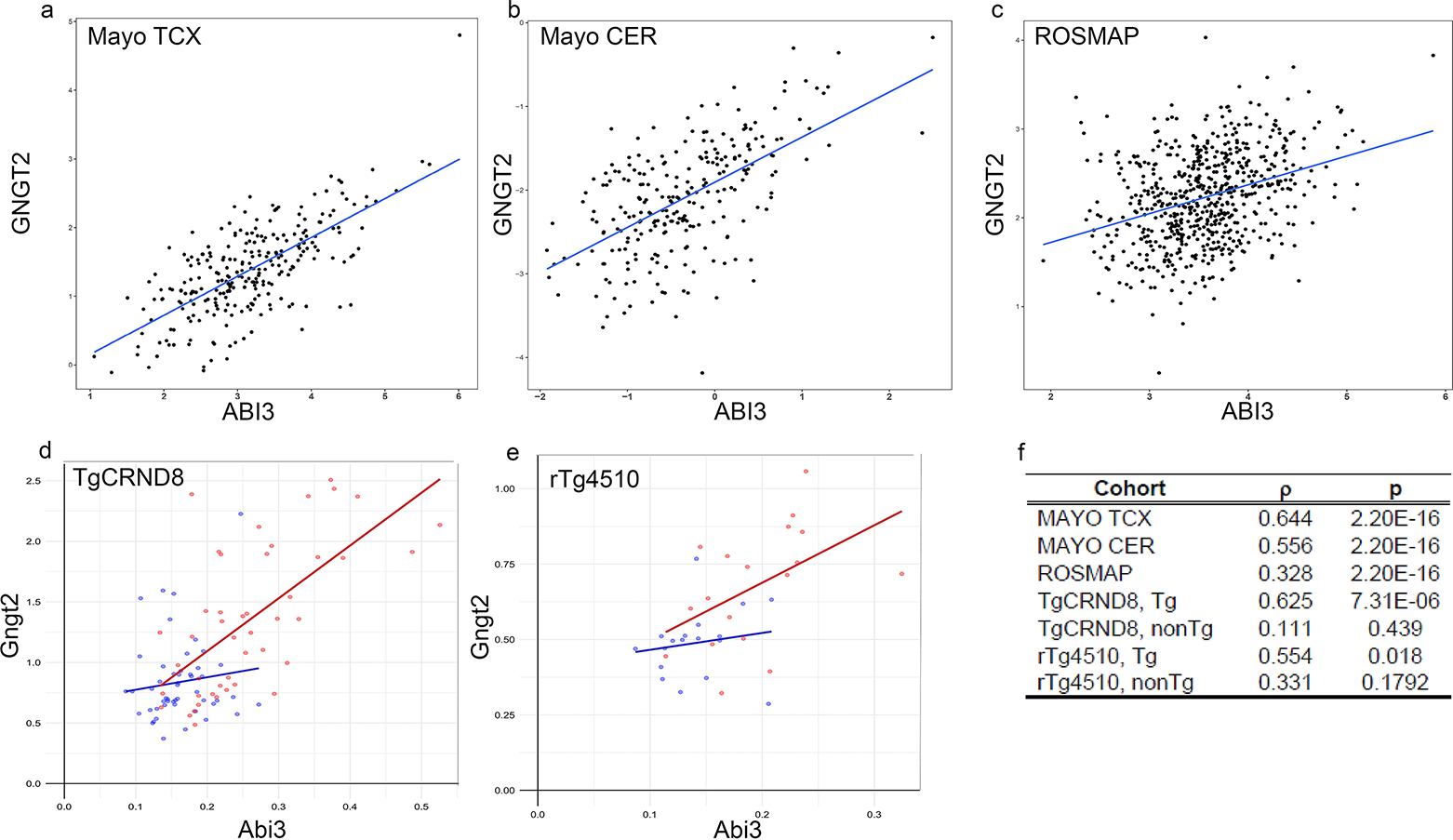
ABI3 and GNGT2 genes are in a co-regulatory expression network. a-c. Graphs showing co-regulation of ABI3 and GNGT2 RNA counts from Mayo cohorts (temporal cortex, TCX and Cerebellum, CER) and ROSMAP cohort. Data derived from AMP-AD synapse. d-e. Graphs showing co-regulation of ABI3 and GNGT2 RNA counts from *APP* TgCRND8 and *MAPT* rTg4510 mice. ρ indicates correlation coefficient and p values are adjusted for multiple analysis. Data derived from AMP-AD synapse. f. Table depicting the correlation coefficient (ρ) and p values for each cohort depicted in a-e.

### Loss of *Abi3*-*Gngt2* induces reactive gliosis and a glial gene signature typically associated with neurodegenerative proteinopathy

We evaluated baseline gliosis in the parental *Abi3-Gngt2*^-/-^ line. Using Iba-1 immunohistochemistry, we found that at 3 months of age, the haploinsufficient *Abi3-Gngt*^+/-^ mice showed reduced hippocampal microgliosis compared to mice with intact *Abi3-Gngt2^+/+^* alleles (p<0.01 in hippocampus) and *Abi3-Gngt2*^-/-^ mice (ns association) (Fig. 3a-c). At 6 months of age, we noticed an interesting gene-dosage dependent dichotomy in microgliosis in the parental *Abi3-Gngt2*^+/-^ lines. The haplo-sufficient *Abi3-Gngt2*^+/-^ showed higher Iba-1 reactivity relative to *Abi3-Gngt2^+/+^* mice (p<0.05 in cortex) and *Abi3-Gngt2*^-/-^ mice (p<0.01 in cortex and p<0.05 in hippocampus) (Fig. 3d-f). There were no significant differences in microgliosis between the *Abi3-Gngt2^+/+^* and *Abi3-Gngt2*^-/-^ mice at this age (Fig. 3d-f).

**Fig. 3.**
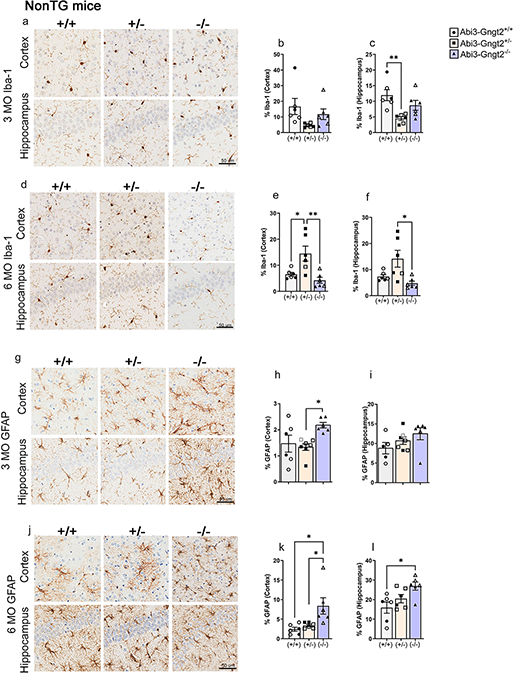
Immune activation in *Abi3*-*Gngt2* deficient mice. Representative images of Iba-1 reactive microglia (a-f) and GFAP reactive astrocyte (g-l) in 3 month old or 6 month old mice with intact (+/+), partial loss (+/-) or complete loss (-/-) of *Abi3-Gngt2* genes. Quantitation of the Iba-1 or GFAP staining from cortex or hippocampus is provided in corresponding panels on the right side. N=6 mice (a-f), 7 mice (g-l). Scale bar, 50µm. Clear symbols indicate female mice and filled symbols represent male mice. n=4 females, 2 males. Data represents mean±sem. 1 way Anova; ***p<0.001; **p<0.01; *p<0.05.

At 3 months of age, the parental *Abi3-Gngt2*^-/-^ line had significantly higher astrocyte burden than the *Abi3-Gngt2*^+/+^ (p<0.05 in cortex and hippocampus) or *Abi3-Gngt2*^+/-^ mice (p<0.01 in cortex) (Fig. 3g-i). At 6 months of age, the astrocyte burden continued to remain elevated in the *Abi3-Gngt2*^-/-^ mice relative to *Abi3-Gngt2^+/+^*mice (p<0.05 in cortex and hippocampus) and *Abi3-Gngt2*^+/-^ mice (p<0.05 in hippocampus) (Fig. 3j-l). This shows that complete loss of function of the *Abi3* locus results in early astrocytosis that persists as the mice age.

We performed bulk RNAseq from the forebrains of 3 month old *Abi3-Gngt2*^-/-^ mice (Fig. 4; **Fig. EV3**). The upregulated RNAs in the *Abi3-Gngt2*^-/-^ mice were predominantly microglia-specific, such as C-type lectin domain family 7 member A (Clec7a/Dectin 1), Macrophage Expressed 1 (Mpeg1), Natural resistance-associated macrophage protein 1 (Slc11a1/Nramp1) and Lymphocyte Antigen 86 (Ly86) and Olfactomedin-like 3 (Olfm13) (Fig. 4a-b). Downregulated RNAs included Abi3, Gngt2, G Protein-Coupled Receptor 179 (Gpr179) and TNF Receptor Superfamily Member 1B (Tnfrsf1b) (Fig. 4a-b). Analysis of the *Abi3-Gngt2*^+/-^ mice relative to *Abi3-Gngt2^+/+^* mice revealed only one significantly altered (downregulated) gene -Protocadherin Gamma Subfamily A 5 (Pcdha5) - that is involved in establishing and maintaining intercellular connections in the brain (**Fig. EV3a**). Pathway analysis of the differentially expressed gene sets in *Abi3-Gngt2*^-/-^ mice vs *Abi3-Gngt2^+/+^* mice revealed the involvement of immune pathways, such as granulocyte (GO:0071621) and leukocyte chemotaxis (GO:0030595), proliferation of mononuclear leukocytes (GO:0032943) and leukocyte-mediated immunity (GO:0002443) (Fig. 4c). The cell types most affected in *Abi3-Gngt2*^-/-^ mice were microglia (p<0.05 relative to both *Abi3-Gngt2^+/+^* and *Abi3-Gngt2*^+/-^) (Fig. 4d). There was a specific reduction in neuronal gene counts in *Abi3-Gngt2*^-/-^ mice (p<0.01 relative to *Abi3-Gngt2^+/+^*) and in *Abi3-Gngt*^+/-^ mice (p<0.001 relative to both *Abi3-Gngt2^+/+^* and *Abi3-Gngt2*^-/-^) (Fig. 4f). No significant changes in astrocyte or oligodendrocyte-specific gene expression were seen among the three groups (Fig. 4e**, g**). Surprisingly, the *Abi3-Gngt2*^-/-^ mice by itself showed induction of the amyloid/AD-associated PIG network, even in the absence of Aβ (p<0.01 relative to both *Abi3-Gngt2^+/+^* and *Abi3-Gngt2*^+/-^ mice) (Fig. 4h). We also observed suggestive upregulated trends in the DAM, MGnD, ARM and A1 co-expression networks in the *Abi3-Gngt2*^-/-^ mice (Fig. 4i-l). Weighted gene co-expression network analysis (WGCNA) identified several gene modules correlating the *Abi3*-*Gngt2* genotype with the gliosis phenotype (Fig. 4n). The hub genes of selected modules that specifically correlated with the *Abi3-Gngt2^-/-^* genotype and gliosis include Unc93b1 (antiquewhite2) and Immunoglobulin kappa variable 10-96 (coral2) (**Fig. EV3b-g**). Genes within the antiquewhite module were previously reported to be associated with gene signatures in AD, most notably DAM and MGnD (Keren-Shaul et al., 2017, Krasemann et al., 2017), PIG ((Deming et al., 2019) and homeostatic microglia (Krasemann et al., 2017) (**Fig. EV3h**). The antiquewhite2 module is especially relevant to AD pathophysiology as the module members, Ctss, Siglec h, Csf3r, Ly86 and C1qc, have been reported in both mouse models and humans (Walker, 2020). It was also highly associated with several immune and autoimmune conditions such as Staphylococcus infection and primary immunodeficiency as well as neurodegenerative diseases such as prion disease (Fig. 4o). Overall, RNA-seq data shows upregulation of an immune activation gene expression profile corresponding to AD-typical gene expression patterns in *Abi3*-*Gngt2*^-/-^ mice, even in the absence of Aβ.

**Fig. 4.**
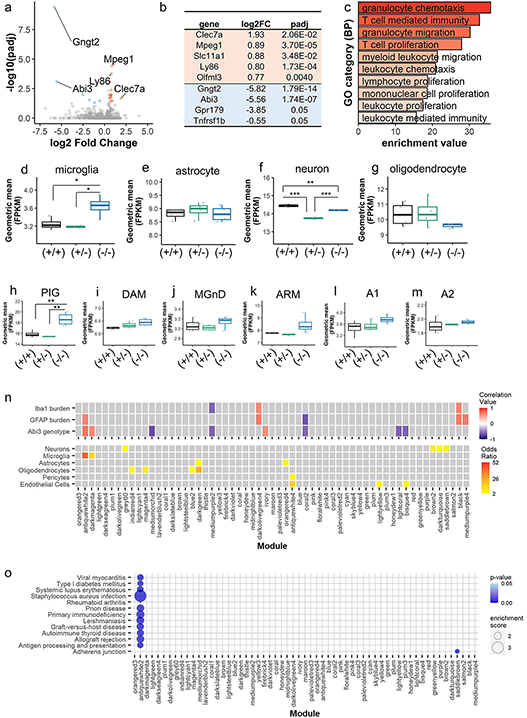
Unbiased transcriptomic analysis of *Abi3*-*Gngt2* deficient mice reveal upregulation of immune pathways and disease-related gene expression patterns. a-c. Volcano plot (a), list of top 5 upregulated and top 5 downregulated genes (based on fold change) (b) and GO pathways based on enriched genes (c) in 3 month old *Abi3-Gngt2*^-/-^ mice relative to *Abi3-Gngt2*^+/+^ mice. Orange dots, upregulated genes; blue dots, downregulated genes. FC= fold change. d-g. Gene expression signatures of microglia (d), astrocytes (e), neurons (f) and oligodendrocytes (g) in 3 month old mice with intact (+/+), partial loss (+/-) or complete loss (-/-) of *Abi3-Gngt2* locus. h-m. Gene expression signatures for specific microglia (h-k) or astrocyte (l, m) subtypes in 3 month old mice with intact (+/+), partial loss (+/-) or complete loss (-/-) of *Abi3*-*Gngt2* locus. These cell-type specific signatures were identified in previous studies (Chen et al., 2020, Keren-Shaul et al., 2017, Krasemann et al., 2017, Liddelow et al., 2017, Sala Frigerio et al., 2019). n-o. WGCNA gene co-expression modules correlating with specific experimental traits (inflammation burden and *Abi3*-*Gngt2*^-/-^ genotype) shown (n). Correlation of modules to different experimental traits are colored in a heatmap (red, positive correlation; blue, negative correlation). Modules with p-values<0.05 and an absolute correlation value < 0.5 are indicated in grey. Cell-type specific gene lists were used to identify genes with significant overlap (odds ratio) within the modules. The heatmap is colored by the value of the odds ratio. Grey squares indicate non-significant (p<0.05, odds ratio < 2) overlaps in the gene lists. o. Genes within WGCNA modules enriched for known KEGG pathways based on over-representation of enriched genes is depicted in a bubble plot (o). Pathways with an over-represented p-value ≤ 0.05, the number of module genes within the pathway >5 and an enrichment score >1.5 are depicted. The bubble plot is colored by p-value and sized by the enrichment score. All p values adjusted for multiple comparisons (padj). n=4 mice (2 males, 2 females) per *Abi3-Gngt2* genotype except 1 outlier removed in d-m. 1 way Anova (D-M), **p<0.01, *p<0.05.

### Reduction in Aβ levels in *APP* mice lacking *Abi3-Gngt2*

We then examined how complete insufficiency or haploinsufficiency of the *Abi3* locus alters development of Aβ plaques in *APP* transgenic CRND8 mice. Transgenic *APP* mice (referred as ‘TG’) that are wild type (+/+), haploinsufficient (+/-) or knocked out (-/-) for *Abi3*-*Gngt2* genes were aged to 3 and 6 months and the Aβ levels assessed by immunohistochemistry and biochemical analysis (Fig. 5, **Fig. EV4**). There was no change in APP expression levels in these bigenic TG-*Abi3-Gngt2* colonies (**Fig. EV4a-b**). In 3 month old TG-*Abi3/Gngt2* lines, both *Abi3-Gngt2*^+/-^ (p<0.05) and *Abi3-Gngt2*^-/-^ (p<0.01) mice showed reduced Aβ plaques relative to littermate *APP* transgenic mice wild type for *Abi3-Gngt2^+/+^* (Fig. 5a-b). Concurrently, there was a non-significant reduction in the number of Thioflavin S cored plaques in the TG-*Abi3-Gngt2*^-/-^ mice (Fig. 5c-d). Biochemical analysis showed significant reduction of FA associated insoluble Aβ42 and Aβ40 levels in both TG-*Abi3-Gngt2*^+/-^ (Aβ42: p<0.01; Aβ40: p<0.05) and TG-*Abi3-Gngt2*^-/-^ (Aβ42: p<0.001; Aβ40: p<0.05) mice relative to littermate TG-*Abi3-Gngt2^+/+^* mice (Fig. 5e-f). In the SDS detergent soluble fraction, both Aβ42 and Aβ40 values were reduced in TG-*Abi3-Gngt2*^-/-^ mice (p<0.05) while only Aβ42 was significantly reduced in TG-*Abi3-Gngt2*^+/-^ mice (p<0.05) relative to littermate TG-*Abi3-Gngt2^+/+^* mice (Fig. 5g-h). We did not observe major changes in ubiquitin labeling of the Aβ plaques across all the *Abi3* genotypes (**Fig. EV4c**).

In 6 month old TG-*Abi3-Gngt2* lines, complete deletion of the locus resulted in reduced total Aβ plaque burden (Fig. 5i-j, p<0.05) as well as number of cored plaques (Fig. 5k-l, p<0.01). Notably, though the immunohistochemical plaque burden was similar between TG-*Abi3-Gngt2*^+/-^ and TG-*Abi3-Gngt2^+/+^* mice (Fig. 5i-j), the former group showed lower thioflavin S (ThioS) plaque number compared to the latter group (Fig. 5k-l, p<0.05). The FA and SDS level of Aβ was mostly equivalent among all the groups except for reduction in SDS-associated Aβ40 in TG-*Abi3-Gngt2*^-/-^ mice relative to TG-*Abi3-Gngt2^+/+^* mice (Fig. 5m-p, p<0.05). The patterns of ubiquitin staining around plaques appeared unchanged across the *Abi3* genotypes (**Fig. EV4d**).

**Fig. 5.**
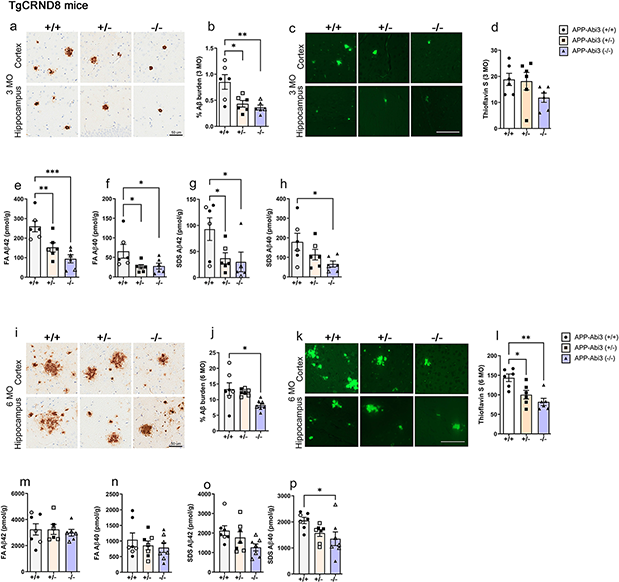
Loss of Abi3-Gngt2 expression ameliorates Aβ in a gene-dosage manner. a-b. Representative immunohistochemical images of Aβ plaques and quantification in 3 month old *APP* TG mice with intact (+/+), partial loss (+/-) or complete loss (-/-) of *Abi3-Gngt2* locus. c-d. Representative images of thioflavin S-stained Aβ plaques and quantitation in 3 month old *APP* TG mice with intact (+/+), partial loss (+/-) or complete loss (-/-) of *Abi3-Gngt2* locus. e-h. Biochemical levels of formic acid (FA) solubilized and detergent (SDS) soluble Aβ42 and Aβ40 in 3 month old *APP* TG mice with intact (+/+), partial loss (+/-) or complete loss (-/-) of *Abi3-Gngt2* locus. i-j. Representative immunohistochemical images of Aβ plaques and quantification in 6 month old *APP* transgenic mice with intact (+/+), partial loss (+/-) or complete loss (-/-) of *Abi3-Gngt2* locus. k-l. Representative images of thioflavin S-stained plaques and quantitation in 6 month old *APP* TG mice with intact (+/+), partial loss (+/-) or complete loss (-/-) of *Abi3-Gngt2* locus. m-p. Biochemical levels of FA solubilized and SDS soluble Aβ42 and Aβ40 in 6 month old *APP* TG mice with intact (+/+), partial loss (+/-) or complete loss (-/-) of *Abi3-Gngt2* locus. N=6 mice (a-h), 7 mice (i-p). Scale bar, 50µm (a, k); 100µm (c, m). Clear symbols indicate female mice and filled symbols represent male mice. n=3 females, 3 males (3 months) and n=3 females, 4 males (6 months). Data represents mean±sem. 1 way Anova; ***p<0.001; **p<0.01; *p<0.05.

We generated an additional cohort of 9 month old TG-*Abi3-Gngt2^+/-^* mice for neuropathological comparisons with age-matched TG-*Abi3-Gngt2^+/+^* mice. We found no changes in Aβ burden, astrocytosis (GFAP), microgliosis (Iba-1 and cd11b) or ubiquitin staining patterns between these two cohorts of mice at this age (**Fig. EV4e-m**).

Alterations in gliosis, especially astrocyte dysfunction, impacts synaptic health (Liu et al., 2018). To survey how amyloid plaques and gliosis in this model affect neuronal health, we evaluate several pre- and post-synaptic proteins in the 3 month time point. Consistent with the reduction in Aβ levels, we saw improved synaptic function as exemplified by increased synaptophysin level in both TG-*Abi3-Gngt2*^+/-^ mice (p<0.05) and TG-*Abi3-Gngt2*^-/-^ mice (P<0.01) (**Fig. EV5a-b**). We observed an insignificant trend in increased PSD95 levels in TG-*Abi3-Gngt2*^-/-^mice while levels of synaptogyrin3 and spinophilin were unaltered in the three genotypes rested (**Fig. EV5c-h**). There was a reduction in vGlut1 (p<0.01 in TG-*Abi3-Gngt2*^+/-^; p<0.05 in TG-*Abi3-Gngt2*^-/-^), GluR1 (p<0.05 in TG-*Abi3-Gngt2*^+/-^) and GluR2 levels (ns trend in TG-*Abi3-Gngt2*^-/-^), signifying dysfunctional glutamatergic signaling (**Fig. EV5i-n**).

### Immunohistochemical and RNAseq shows gene dose-dependent upregulation of inflammatory profile in TG-*Abi3*-*Gngt2^-/-^* mice

Aβ levels are generally well-correlated with immune activation indicated by microgliosis and astrocytosis (Kinney et al., 2018). Consistent with this idea, we predicted lower burden of microglia (Iba-1 immunoreactivity) and astrocytes (GFAP immunoreactivity) in TG-*Abi3-Gngt2*^-/-^ mice that showed robust Aβ reduction at 3 months and 6 months of age. Surprisingly, we found that at both ages, the TG-*Abi3-Gngt2^-/-^* mice showed similar levels of Iba-1 reactive microgliosis compared to TG-*Abi3-Gngt2^+/+^* mice (3 months: Fig. 6a-c; 6 months: Fig. 6d-f). TG-*Abi3-Gngt2^+/-^* mice showed decreased cortical microglia at 3 months compared to TG-*Abi3-Gngt2^-/-^* mice (Fig. 6a-c; p<0.05), but this normalized to equivalent levels by 6 months of age (Fig. 6d-f). The patterns of astrocyte burden reflected a differential scenario across the three *Abi3* genotypes. At 3 months of age, the TG-*Abi3-Gngt2^+/-^* mice had significantly lower astrocytosis compared to both TG-*Abi3-Gngt^+/+^* (p<0.01 in cortex; p<0.001 in hippocampus) and TG-*Abi3-Gngt2^-/-^* mice (p<0.001 in cortex; p<0.01 in hippocampus) (Fig. 6g-i). At 6 months of age, these differences normalized in the cortex but hippocampal GFAP burden in TG-*Abi3-Gngt2^+/-^* mice was higher than both TG-*Abi3-Gngt^+/+^* or TG-*Abi3-Gngt2^-/-^* mice (Fig. 6j-l). There was no significant difference in astrocytosis burden between the TG-*Abi3-Gngt^+/+^* or TG-*Abi3-Gngt2^-/-^* mice in 3 month or 6 month old cohorts (Fig. 6g-i, **6j-l**), indicating that reduction of Aβ did not ameliorate the existing astrocytic phenotype inherent in the *Abi3*-*Gngt2* line. Overall, the immunohistochemistry data suggests a biphasic age-dependent response of the astrocytes and microglia in *APP* mice to *Abi3-Gngt2* gene dosage.

**Fig. 6.**
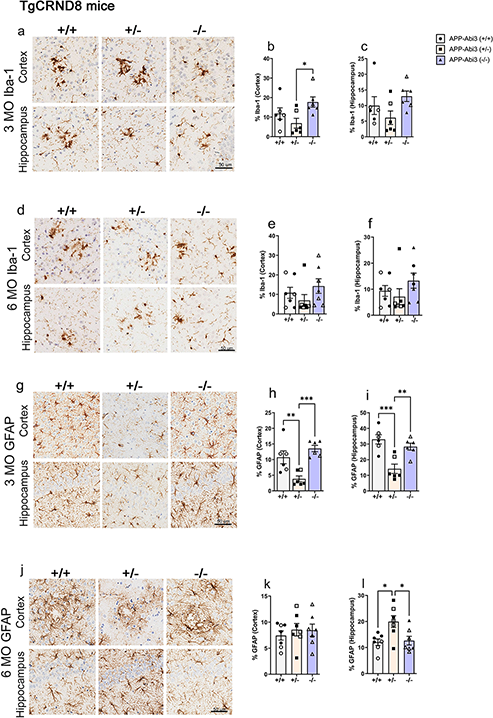
Abi3-Gngt2 regulates microgliosis and astrocytosis in *APP* mice. Representative images of Iba-1 reactive microglia (a-f) and GFAP reactive astrocyte (g-l) in 3 month old or 6 month old *APP* transgenic mice with intact (+/+), partial loss (+/-) or complete loss (-/-) of *Abi3* locus. Quantitation of the Iba-1 or GFAP staining from cortex or hippocampus is provided in corresponding panels on the right side. Scale bar, 50µm. Clear symbols indicate female mice and filled symbols represent male mice. n=3 females, 3 males (3 months) and n=3 females, 4 males (6 months). Data represents mean±sem. 1 way Anova; ***p<0.001; **p<0.01; *p<0.05.

We performed bulk RNAseq from the forebrains of 3 month old TG-*Abi3-Gngt2* mice (Fig. 7, **Fig. EV6**). Relative to the *TG-Abi3-Gngt^+/+^* mice, the *TG-Abi3-Gngt2^-/-^* mice showed lower Abi3 and Gngt2 as expected (Fig. 7a-b). Other genes that were down-regulated in the *TG-Abi3-Gngt2^-/-^* mice are *Adgrf3* (Adhesion G Protein-Coupled Receptor F3), *S100a8* and *S100a9* (S100 Calcium Binding Protein members A8 and A9). Among the genes that were upregulated in these mice were *Gm7879* (Peptidylprolyl isomerase H), *Aklr1c13* (Aldo-keto reductase family 1 member C13), *Fndc9* (Fibronectin Type III Domain Containing 9), tetraspanin *Cd9* and Lymphocyte antigen *Ly86* (Fig. 7a-b). The molecular pathways represented by gene expression changes in *TG-Abi3-Gngt2^-/-^* mice include syncytium formation, cellular fusion, calcium mediated signaling and extracellular matrix organization (Fig. 7c), recapitulating expected functional properties of the ABI family members (Mendoza, 2013). Additional pathways that were enriched were gliogenesis (GO:0014015) and response to LPS (GO:0034189), consistent with altered glial homeostasis. We did not observe any significant gene expression changes in *TG-Abi3-Gngt2^+/-^* mice relative to *TG-Abi3-Gngt^+/+^*mice (**Fig. EV6a**). In the *TG-Abi3-Gngt2^-/-^* mice, most of the gene expression changes were indicative of increased microglial (p<0.05 relative to *TG-Abi3-Gngt2^+/-^*) and astrocytic involvement (p<0.01 relative to *TG-Abi3-Gngt2^+/-^*; p<0.05 relative to *TG-Abi3-Gngt^+/+^*) (Fig. 7d-e), with no changes observed in neuronal and oligodendrocyte-specific gene expression (Fig. 7f-g). Surprisingly, we found that in spite of reduced Aβ plaques, *TG-Abi3-Gngt2^-/-^* mice showed gene signatures typically identified in AD tissues or preclinical models of AD. These mice showed upregulated PIG (p<0.05 relative to *TG-Abi3-Gngt2^+/-^*; p<0.05 relative to *TG-Abi3-Gngt^+/+^*) (Chen et al., 2020), DAM (p<0.05 relative to *TG-Abi3-Gngt2^+/-^*) (Keren-Shaul et al., 2017), MGnD (p<0.05 relative to *TG-Abi3-Gngt2^+/-^*) (Krasemann et al., 2017) and A1 astrocyte (p<0.05 relative to *TG-Abi3-Gngt^+/+^*) (Liddelow et al., 2017) gene profile signatures (Fig. 7h, i, j, l). We did not detect any selective induction of either the ARM (Sala Frigerio et al., 2019) or A2 astrocyte (Liddelow et al., 2017) phenotypes in the three TG-*Abi3/Gngt2* genotypes (Fig. 7k,m). Notably, most of these modules (PIG, DAM and MGnD) are driven by Apoe, Tyrobp and Trem2 while the ARM signature is driven by specialized microglial subgroups overexpressing MHC type II genes. WGCNA identified several gene co-expression modules that allowed us to correlate neuropathological traits to cell types (Fig. 7n), GO pathways (Fig. 7o), hub genes (**Suppl. Fig. EV6b-g**) and AD gene expression signatures (**Suppl. Fig. EV6h**). Modules that were positively correlated with the *Abi3-Gngt2* genotype but were negatively correlated with Aβ biochemical levels or Aβ plaque burden included honeydew1 and plum3. These gene modules were primarily driven by microglia- and astrocyte-specific genes, respectively. We identified hub genes that regulate these different gene network modules (**Fig. EV6b-e**). Notably, Chil1/Chi3l1/YKL-40 (Chitinase-like 1) that is the top hub gene in honeydew1 is related to inflammation and AD pathophysiology (Llorens et al., 2017). The top hub gene in the plum3 module – Interferon Regulatory Factor 8 (Irf8) – correspond to interferon signaling that has recently been identified to be upregulated in microglia from human AD (Zhou et al., 2020). This module is especially enriched for immune function, with the cd37 hub gene related to a Tyrobp-regulated microglial module in AD (Zhangi et al., 2013) and Lat2 identified as a core transcriptional signature of AD microglia (Patir et al., 2019). Conversely, modules negatively correlated with the *Abi3-Gngt2* genotype but were positively correlated with Aβ biochemical levels or Aβ plaque burden was sienna3 which included a mixture of astrocyte and endothelial genes (Fig. 7n**, Fig. EV6f-g**). Among these co-expression modules, we found that the plum3 module corresponds to homeostatic microglia signature (**Fig. EV6h**). Overall, RNAseq and immunohistochemical data shows both amyloid-independent and amyloid-dependent dysfunctional immune signature in the *Abi3-Gngt2*^-/-^ mice.

**Fig. 7.**
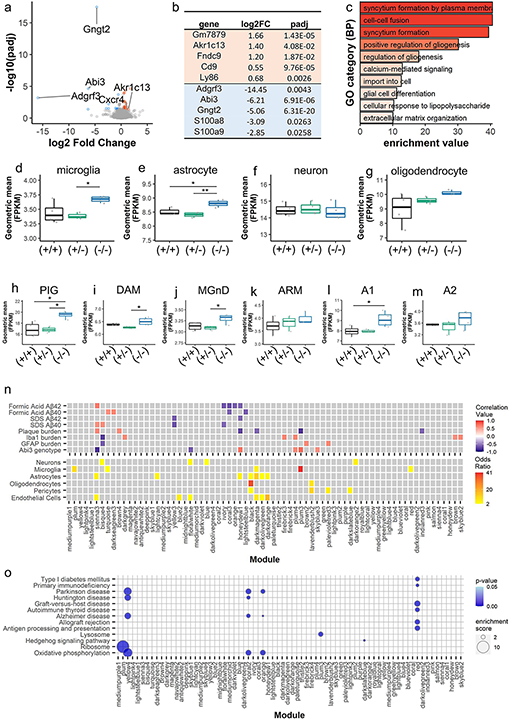
Unbiased transcriptomic analysis of TG-*Abi3-Gngt2^-/-^* mice reveal distinctive disease-associated gene expression signatures and co-expression modules. a-c. Volcano plot (a), list of top 5 upregulated and top 5 downregulated genes (based on fold change) (b) and GO pathways based on enriched genes for upregulated genes (c) in 3 month old *APP* TG mice with intact (+/+), partial loss (+/-) or complete loss (-/-) of *Abi3-Gngt2* locus. Orange dots, upregulated genes; blue dots, downregulated genes. FC=fold change. d-g. Gene expression signatures of microglia (d), astrocytes (e), neurons (f) and oligodendrocytes (g) in 3 month old *APP* TG mice with intact (+/+), partial loss (+/-) or complete loss (-/-) of *Abi3-Gngt2* locus. h-m. Gene expression signatures for specific microglia (h-k) or astrocyte (l, m) subtypes in 3 month old *APP* TG mice with intact (+/+), partial loss (+/-) or complete loss (-/-) of *Abi3-Gngt2* locus. n-o. WGCNA gene co-expression modules correlating with experimental traits (Aβ values, inflammation burden) with *Abi3*-*Gngt2* genotype shown (n). Correlation of modules to different experimental traits are colored in a heatmap (red, positive correlation; blue, negative correlation). Modules with p-values<0.05 and an absolute correlation value < 0.5 are indicated in grey. Cell-type specific gene lists were used to identify genes with significant overlap (odds ratio) within the modules. The heatmap is colored by the value of the odds ratio. Grey squares indicate non-significant (p<0.05, odds ratio < 2) overlaps in the gene lists. o. Genes within WGCNA modules enriched for known KEGG pathways depicted in a bubble plot. Pathways with an over-represented p-value ≤ 0.05, the number of module genes within the pathway >5 and an enrichment score >1.5 are depicted. The bubble plot is colored by p-value and sized by the enrichment score. N=4 mice (2 males, 2 females) per *Abi3-Gngt2* genotype except 1 outlier removed in d-m. All p values adjusted for multiple comparisons (padj). 1 way Anova (D-M); **p<0.01, *p<0.05.

### Exacerbated ptau accumulation in *Abi3-Gngt2^-/-^* mice expressing human mutant tau

Because of the inherent dysfunctional immune milieu and early astrocytosis in the parental *Abi3-Gngt2^-/-^* mice, we decided to test how this would affect the development and progression of tauopathy. We used recombinant AAVs to deliver the pro-aggregant P301L/S320F human tau, WT human tau or a control vector in neonatal *Abi3-Gngt2^-/-^* mice or *Abi3-Gngt2^+/+^* C57BL/6 mice (Beckman et al., 2021, Koller, 2019) (Fig. 8; **Fig. EV7**). We examined mice at 3 months of age and 6 months of age. We first confirmed that the expression of tau transgene was equivalent in P301L/S320F tau expressing *Abi3-Gngt2^-/-^* and *Abi3-Gngt2^+/+^* mice at 3 months (Fig. 8a,b) and 6 months (Fig. 8g,h). We did not find any human tau signal in the control vector injected *Abi3-Gngt2^-/-^* mice or *Abi3-Gngt2^+/+^* mice as expected (Fig. 8a,b,g,h). We noticed that the level of phosphorylated tau (ptau) and misfolded pretangle tau was significantly higher in 3 month old P301L/S320F tau expressing *Abi3-Gngt2^-/-^* mice compared to *Abi3-Gngt2^+/+^*mice (p<0.05 for CP13 and MC1 respectively) (Fig. 8c,d,e,f). None of the control injected mice showed any detectable ptau or misfolded tau at either age (Fig. 8c-f, i-l). We also confirmed the presence of frank neurofibrillary tangles (NFT) in P301L/S320F tau expressing 3 month old *Abi3-Gngt2^-/-^* and *Abi3-Gngt2^+/+^* mice by staining with ThioS (**Fig. EV7a**) or by biochemical analysis of insoluble tau following sequential extraction of cell lysates (**Fig. EV7b-c**). At 6 months of age, the P301L/S320F tau expressing *Abi3-Gngt2^-/-^* mice showed higher ptau immunoreactivity compared to *Abi3-Gngt2^+/+^* mice (Fig. 8i-j; p<0.001), while the levels of pretangle tau were equivalent between the two groups (Fig. 8k-l).

**Fig. 8.**
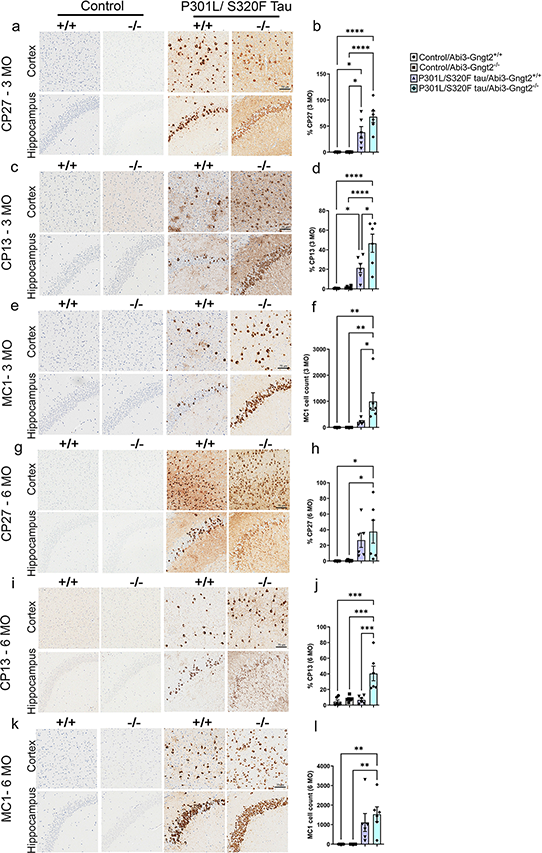
Abi3-Gngt2 deficiency accelerates tauopathy in a mutant human tau model. Mice were injected with control vector (Control) or AAV expressing a double mutant (P301L/S320F) human tau on neonatal day P0 and analyzed at 3 months (a-f) or 6 months (g-l). Representative brain images of total human tau (detected with CP27 antibody, a, g), ptau (detected with CP13 antibody, c, i) and misfolded pretangle tau (detected with MC1 antibody, e, k) from cortex and hippocampus of wild type BL/6 mice (+/+) and *Abi3*-*Gngt2*^-/-^ (-/-) mice are shown. Quantitative analysis of antibody-stained slides (b, d, f, h, j, l) shown on right side of corresponding stained brain images (a, c, e, g, i, k). Scale bar, 75 µm. n=6 mice/group. Data represents mean±sem. 1 way Anova; ****p<0.0001; ***p<0.001; **p<0.01; *p<0.05.

We also examined the neuropathologies in human WT tau expressing *Abi3-Gngt2^-/-^* mice and *Abi3-Gngt2^+/+^* mice. In the 3 month old cohorts, we did not find any differences in the total tau levels or ptau levels of *Abi3-Gngt2^-/-^* and *Abi3-Gngt2^+/+^* mice (**Fig. EV7d-g**), nor did we see any induction of MC1-reactive pretangle tau (data not shown). In the 6 month old cohort, while the levels of tau expression were comparable (**Fig. EV7l-m**), we noticed considerably higher ptau levels in *Abi3-Gngt2^-/-^* mice relative to *Abi3-Gngt2^+/+^* mice (p<0.0001) (**Fig. EV7n-o**). Overall, loss of *Abi3*-*Gngt2* resulted in increased ptau in two separate tau cohorts.

### Astrocyte activation following tau overexpression in *Abi3-Gngt2^-/-^* mice

We wanted to examine whether the inherent immune phenotype in *Abi3-Gngt2^-/-^* mice would be exacerbated in the presence of ptau (Fig. 9). In 3 month old mice, P301L/S320F tau expression had a robust microglial response in the *Abi3-Gngt^+/+^* mice (p<0.01 relative to control vector; p<0.05 relative to tau expressing *Abi3-Gngt2^-/-^* mice) (Fig. 9a-b). At 6 months of age, the P301L/S320F tau expressing *Abi3-Gngt2^-/-^* mice showed higher microglial response than the rest of the cohorts (p<0.01 relative to control vector) (Fig. 9c-d). In WT tau expressing mice, we observed an identical age-dependent pattern for microgliosis with highest levels of microgliosis in 6 month old *Abi3-Gngt2^-/-^* mice (**Fig. EV7j, k, r, s).**

**Fig. 9.**
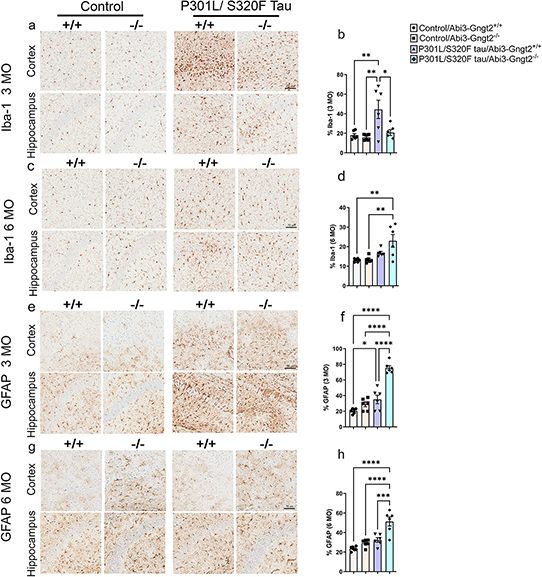
Upregulation of astrocytosis in tau expressing *Abi3-Gngt2^-/-^* mice. Mice were injected with control vector (Control) or AAV expressing a double mutant (P301L/S320F) human tau on neonatal day P0 and analyzed at indicated time points. Representative images of Iba-1 reactive microglia (a-d) and GFAP reactive astrocyte (e-h) in 3 month old or 6 month old mice with intact (+/+) or complete loss (-/-) of *Abi3-Gngt2* locus. Quantitation of the Iba-1 or GFAP staining is provided in corresponding panels on the right side. N=6 mice/group. Scale bar, 75 µm. Data represents mean±sem. 1 way Anova; ****p<0.0001; ***p<0.001; **p<0.01; *p<0.05.

In the 3 month old and 6 month old P301L/S320F tau expressing *Abi3-Gngt2^-/-^* mice, we observed higher astrocytosis concordant with increased pathological tau accumulation (Fig. 9e-h). P301L/S320F tau expressing *Abi3-Gngt2^-/-^* showed higher GFAP burden relative to control vector (p<0.0001) as well as P301L/S320F tau expressing *Abi3-Gngt2^+/+^* mice (p<0.001) at both ages. Thus, the astrocytic response was more consistent with the neuropathology at both ages. This data also reveals an interesting dichotomy in the response of astrocytes and microglia to pathological tau, especially at the younger age examined (Fig. 9b vs Fig. 9f). In 3 month old WT tau expressing mice, astrocytosis immunoreactivity was more enhanced in *Abi3-Gngt2^+/+^* mice than *Abi3-Gngt2^-/-^* mice (p<0.05) (**Fig. EV7h-i**). At 6 months of age, however, we observed higher levels of astrocytosis in WT tau expressing *Abi3-Gngt2^-/-^* mice compared to *Abi3-Gngt2^+/+^* mice (**Fig. EV7p-q**; p<0.01). This implies an age-dependent as well as neuropathology dependent immune phenotype induced by tau overexpression in the *Abi3-Gngt2^-/-^* mice. Overall, the effect of tauopathy on astrogliosis was most prominent in 6 month old *Abi3-Gngt2^-/-^* mice.

### AD-associated mutation alters ABI3 phosphorylation

To understand the differences between wild type and AD-associated S209F ABI3, we overexpressed C terminal FLAG tagged recombinant constructs in HEK293T cells. C terminally FLAG tagged ABI3 has been previously shown to be functionally active (Sekino et al., 2015). We detected ABI3 protein using either anti FLAG antibody, an in-house antibody raised against S209 ABI3 or a N terminal ABI3 antibody (Millipore Sigma). We observed that wild type ABI3, but not S209F ABI3 migrated as a double band (Fig. 10a) which is consistent with data reported earlier (Sekino et al., 2015). To confirm that the band migrating on the top is a phosphorylated form, we incubated wild type ABI3 or S209F ABI3 transfected HEK293T cell lysates with Lambda Protein Phosphatase. Incubation with Lambda Protein Phosphatase at 30°C but not at 4°C abolished the upper band in wild type ABI3 lysate (Fig. 10b). Mutating the Phenylalanine (F) at position 209 to dephospho-mimetic Alanine (A) or phospho-mimetic Aspartate (D) reverses this deficit in phosphorylation to levels that are comparable to WT ABI3 protein (Fig. 10c-d). This shows that presence of phenylalanine at position 209 specifically impairs overall ABI3 phosphorylation, though auto-phosphorylation at this site is not necessary. Since, phosphorylation of Abi family of proteins is related to their stability and function (Huang et al., 2007), this would imply that the Ser to Phe mutation could render the ABI3 protein functionally inactive.

**Fig. 10.**
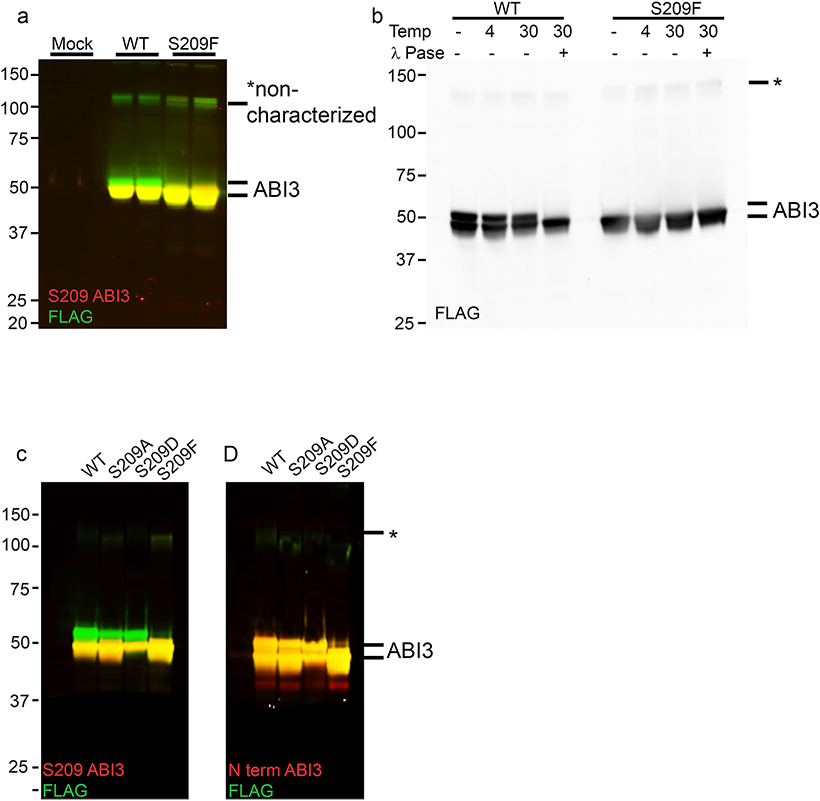
S209F mutation leads to altered phosphorylation in ABI3 protein. a. HEK293T cells were transiently transfected with WT ABI3 or S209F ABI3 and separated on an immunoblot showing two distinct bands around 50kDa. Higher bands that are as yet uncharacterized (marked by asterisk) are also denoted. The proteins were detected with two antibodies simultaneously using LiCor infrared dyes: anti FLAG antibody (green) and anti S209-ABI3 antibody (red). b. Lambda protein phosphatase assay performed at 30°C on WT ABI3 and S209F ABI3 protein detected with anti FLAG antibody. c-d. HEK293T cells were transiently transfected with WT ABI3 or ABI3 mutated at position 209 and separated on an immunoblot. The proteins were detected with two antibodies simultaneously using LiCor infrared dyes: anti FLAG antibody (green) and anti S209-ABI3 antibody (red) or anti N terminal ABI3 (Sigma) (red). ‘Mock’ denotes sham transfection procedure with no DNA used in transfection. Asterisk marks an ABI3 derived band that has not been characterized. Data representative of at least two independent experimental replicates.

## Discussion

The mechanism through which ABI3 alters risk for AD remains uncertain. Here we document that Abi3, though primarily expressed in microglia, is also expressed in other cells in human and mouse brain. Age-associated increases in Abi3 mRNA appear to be largely attributable to local increases in Aβ plaque-associated microglia. Characterization of an *Abi3*^-/-^ mouse strain identified that the overlapping gene Gngt2 is also knocked out. We conducted additional studies to demonstrate that *Abi3* and *Gngt2* expression are tightly co-regulated in both mice and humans. Using this *Abi3*-*Gngt2*^-/-^ mice, we show that loss of the Abi3-Gngt2 expression predisposes the mice to an immune phenotype characterized by reactive gliosis and gene expression patterns resembling an AD-associated profile. Notably in these mice, the AD/neurodegeneration-associated genes Trem2, Plcg2, Tyrobp and Csf1r are upregulated, even in the absence of Aβ. We find that the loss of *Abi3*-*Gngt2* locus has opposing effects on amyloid and tau pathology in mice. Loss of this locus results in a reduction in amyloid pathology and preservation of synaptic markers, though effects on amyloid are diminished as the mice age. In contrast, we noted that loss of *Abi3*-*Gngt2* markedly increases tau pathology (Fig. 11). Because Abi3 is expressed in neurons as well as microglia, it is possible that the effect of *Abi3*-*Gngt2* on Aβ and tau could be modulated by glial as well as neuronal signaling pathways. Our experimental models, whereby we observed functional dichotomy in Abi3-mediated immunoproteostasis in Aβ and tau models, allows us to impute the probable function of the risk variant in influencing AD risk. If, for example, the detrimental effect of S209F Abi3 is mediated by increasing Aβ, then the disease-associated mutation is expected to reinforce additional functional characteristics. On the other hand, if the effect is cell autonomous, leading to deregulation of microglial surveillance activity or exacerbating neuronal tauopathy, then the mutation could be mimicking a loss of function phenotype. Indeed, examples of such context-dependent manifestations can be found in Trem2 models, where depending on the disease stage, neuropathology and gene dosage, Trem2 expression (or lack of) recapitulates a partial loss-function or demonstrates additional pathological characteristics (Bemiller et al., 2017, Lee et al., 2021, Leyns et al., 2017, Sayed et al., 2018).

**Fig. 11.**
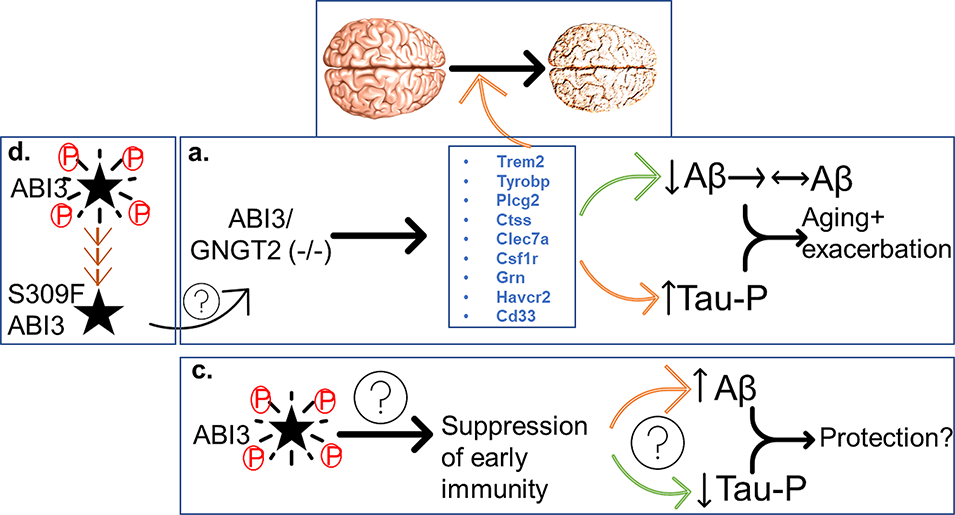
Schematic depiction of Abi3-Gngt2 function. Our experiments with *Abi3*-*Gngt2* deficient model show that immune phenotype in these mice reduce Aβ plaques in a gene-dose and age-dependent manner but increase tau phosphorylation early on (a). Loss of Abi3-Gngt2 function leads to early-life induction of AD/amyloid-associated factors and microglial genetic risk factors, independent of Aβ. In fact, these same microglial factors (as tabulated in a) are upregulated as humans progress from healthy status to AD (b). Based on our data, we hypothesize that a normally functioning Abi3 would lead to suppression of immunity (by suppressing induction of AD disease microglial risk factor genes), thus increasing Aβ plaques but preventing tau phosphorylation and tangle formation (c). The neuropathology in the *Abi3*-*Gngt2* deficient model is reminiscent of the dichotomy observed in multiple mouse models testing microglial AD risk factors. The differential response of Aβ and tau to immune activation necessitates a cautious foray into immune-targeted therapies. Our in vitro data shows that the AD associated S209F mutation dramatically reduces self-phosphorylation, which guided by previous data from ABI family of proteins, would imply a loss of function phenotype (d). Brain picture adapted from online (CC license, b).

Taken together, our data show that a dysfunctional immune environment accompanying the lack of *Abi3*-*Gngt2* is associated with a dichotomous effect on Aβ and tau proteinopathy. Indeed, manipulation of expression of Trem2, Tyrobp, Cx3cl1/Cx3cr1, IL1β and IFN-γ have all been shown to have opposing effects on amyloid and tau pathologies (Audrain et al., 2019, Chakrabarty et al., 2010, Haure-Mirande et al., 2019, Jay et al., 2015, Lee et al., 2021, Leyns et al., 2017, Li et al., 2015, Meiland et al., 2020, Parhizkar et al., 2019, Sayed et al., 2018). We have previously referred to this multi-directional relationship between immunity and AD-type proteinopathy as immunoproteostasis (Chakrabarty et al., 2015, Golde, 2019). Harnessing the immune system as a potential treatment for AD remains attractive. Our current data, however, highlights the challenges of harnessing immunoproteostasis to treat AD as such therapies could theoretically have opposing effects of AD-triggering neurodegenerative proteinopathy and exacerbate underlying brain organ dysfunction.

A recent study, using a bigenic model of 5xFAD and *Abi3*/*Gngt2*^-/-^ mice, reported increased Aβ plaques at 8 months of age and an inflammatory gene expression signature similar to our data (Karahan et al., 2021). However, this report could not specifically attribute the DAM profile to *Abi3*/*Gngt2* genotype (Karahan et al., 2021). While the changes in Aβ proteinopathy is opposite of what we have observed, this could be due to this group using a single time point and a mouse model expressing both mutant *APP* and *PS1* genes. Whether *Abi3*/*Gngt2* could have an additional interaction with PS1-related phenotype remains to be studied. Research groups have reported disparate observations in mouse models of neurodegenerative proteinopathies that undergo immune manipulations. A recent example is when the profound systemic and neural inflammation inherent in the c9orf72 deficient mice was found to be reversed when maintained in a different mouse facility (Burberry et al., 2020). In fact, immune-related phenotypes, as inherent in this Abi3/Gngt2-/-mice, are extremely sensitive to environmental factors and local immune stressors that can regulate the microbiome (Erny et al., 2015).

Our study highlights the utility of detailed transcriptomic analysis in phenotyping new disease models. During our study, we found that the *Gngt2* gene, whose putative promoter region and a non-coding exon of one transcript isoform overlaps with the *Abi3* locus, is also lost in this velocigene knockout model. Gngt2 is reported to be a microglia-specific gene and is involved in cone phototransduction (Ong et al., 1997). To establish the physiological relevance of this dual knock-out, we used extensive informatics analysis from human and mice showing that ABI3 and GNGT2 are co-regulated, indicating that these two genes exist in a co-expression network. While the original exome analysis data (Sims et al., 2017) did not find any AD risk related to *GNGT2*, a newer study identified GNGT2 as one of the 989 genes that mapped to the 38 genomic loci related to AD risk (Wightman et al., 2021). Future studies should dissect out the specific functions of the genes resident in the *ABI3* locus.

Our study is an exemplar for highlighting how immune function can be differentially related to development and progression of AD-typical neuropathologies. Several seminal studies have shown that microglia-mediated immune activation can exacerbate both Aβ and tau pathologies (Friker et al., 2020, Heneka et al., 2013, Ising et al., 2019). More recently however, immune activation has been shown to have disease stage-dependent outcome. Studies in human patients support this notion of a dichotomous role of inflammation, where data indicates that early inflammation may play a protective role in prodromal AD phase (Fan et al., 2017, Hamelin et al., 2016) but that chronic age-associated inflammation may be overwhelmingly detrimental. This dichotomy of immunity having a beneficial as well as a harmful effect has been shown previously for at least two other immune factors in mice. Hippocampal expression of IL-1β was shown to robustly reduce plaques in the APP/PS1 model and 3xTg-AD model but worsened tauopathy in the 3xTg-AD model (Ghosh et al., 2013, Shaftel et al., 2007). Similarly deficiency of Cx3cr1 reduced amyloid deposition in two different *APP* models but aggravated tau hyperphosphorylation in hTau mice (Bhaskar et al., 2010, Lee et al., 2010). More recently, this dichotomy has been elegantly demonstrated in multiple models of Trem2 and Tyrobp (Audrain et al., 2019, Haure-Mirande et al., 2019, Jay et al., 2015, Lee et al., 2021, Leyns et al., 2017, Meilandt et al., 2020, Parhizkar et al., 2019, Sayed et al., 2018). Our previous data (Chakrabarty et al., 2010, Li et al., 2015), as well as our current data, also indicates that immune function can have distinctive effects on Aβ and tau, a paradigm that we have referred to as immunoproteostasis (Golde, 2019). The outcomes of *Abi3*-*Gngt2* related immunoproteostasis was based on age as also on gene dose, consistent with studies on Trem2 models of AD (Meilandt et al., 2020, Sayed et al., 2018). Combined, our data shows that underlying genetic factors that enhance the reactivity of immune cells might result in early beneficial effects on Aβ but this might adversely affect emergence of tauopathy and age-progressive Aβ pathology. Thus, designing immune-based therapies warrants cautious deliberation with careful consideration of disease stage and immune mediator function.

Understanding how immunoproteostasis is modified by brain neuropathology and age is important as this is expected to modify the efficacy of immune-based therapies. For amyloid pathology, there is data that activated micro/astroglia enhances Aβ phagocytosis (Chakrabarty et al., 2010, Chakrabarty et al., 2010a, Shaftel et al., 2007), which is a desired outcome, but why that glial activation state appears to be associated with enhanced tau pathology remains enigmatic. Given that large number of immune signaling pathways are altered in our current study and in previous studies on AD immunoproteostasis (Heppner et al., 2015), it will be challenging to gain more detailed mechanistic insights. Indeed, it may not be a single pathway or factor that results in these age- and pathology-related dichotomous outcomes but the combined action of many different signaling alterations on Aβ and tau. Given that alterations in immune signaling induced by amyloid deposition are a clear hallmark of AD, such data provide an alternative, but not mutually exclusive, mechanisms for spreading of tau pathology. Indeed, signaling initiated by microglia or astrocytes in response to amyloid deposition could then trigger induction of tau pathology. Such signaling could explain the apparent crosstalk between amyloid and tau (Nisbet et al., 2015), characterized by the unique region-specific spread of tau and could explain the temporal lag between amyloid deposition and tau tangle formation in humans.

Analysis of co-expression networks reveals interesting transcriptomic differences in parental *Abi3-Gngt2*^-/-^ lines and the TG*-Abi3-Gngt2*^-/-^ lines. While the parental *Abi3-Gngt2*^-/-^ lines mostly indicated an abundance of immune signature, the TG-*Abi3-Gngt2*^-/-^mice showed additional signatures consistent with the known functions of the ABI family of proteins (Mendoza, 2013). Consideration of these functions – calcium mobilization, extracellular matrix reorganization and intercellular communication (Siddiqui et al., 2012) -will be important in understanding AD pathogenesis in addition to the involvement of more well-characterized phagocytotic and debris removal functions of microglia. Indeed, a recent study suggests that Abi3-Gngt2^-/-^ microglia exhibits reduced surveillance of brain parenchyma (Simonazzi, et al., 2021). In terms of the hub genes identified in our study, Chi3l1/YKL-40 is singularly important as it is correlated with the *Abi3-Gngt2* genotype in our study. YKL-40 is primarily an astrocytic protein that is elevated in AD and is considered a potential biomarker (Craig-Schapiro et al., 2010). Recently YKL-40 has been shown to have non cell autonomous activity in the brain, affecting microglial function, circadian clock and generally AD progression (Lananna et al., 2020). How Abi3 plays a role in AD neuropathologic cascade by modulating YKL-40 function remains an untested possibility.

ABI3 has been imputed to be of microglial origin (Patir et al., 2019). A previous study has reported clusters of ABI3 immunopositive microglia localized around Aβ deposits in AD patients (Satoh et al., 2017). Our in situ hybridization data indicates that ABI3 is present in both microglia and neurons. We also show that the amount of ABI3 RNA increases in concordance with Aβ deposits. Because Abi3 is present in both neurons and microglia, its role in non-cell autonomous and cell autonomous signaling in AD proteinopathy and disease progression could be equally important. Our unexpected finding of neuronal ABI3 is reminiscent of the observations on PLCG2, which was initially thought to be myeloid cell-specific, but its RNA was also found in neurons and endothelial cells (Magno et al., 2019). PLCG2, like ABI3, was identified as an AD protective factor using exome analysis strategies (Sims et al., 2017). Given that protein levels in any specific cell cannot always be inferred from RNA levels (Liu et al., 2016), some caution is warranted here before confirming our in situ observations by protein detection methods.

The ABI family of proteins, especially ABI-1 and ABI-2, are involved in cytoskeletal reorganization (Mendoza, 2013). The function of ABI3, much less S209F ABI3, in the brain is unknown. Some previous studies have reported ABI3 in microglia with ramified or amoeboid morphology (Patir et al., 2019, Satoh et al., 2017). Another recent paper showed that absence of Abi3 (using the same models as used in this study) alters microglial morphology and curbs its function in homeostatic surveillance (Karahan et al., 2021, Simonazzi et al., 2021). This would be consistent with the previously documented role of ABI3 in regulating cytoskeletal dynamics (Bae et al., 2012b, Ichigotani et al., 2002).

The *Abi3*-*Gngt2*^-/-^ mice also show increased Trem2, Tyrobp and Plcg2, which would be consistent with cooperativity between different AD-associated risk factors. In fact, we observed this early immune activation in the absence of Aβ, which could illuminate how underlying genetic risk factors could cross-talk to prime the aging brain, even before the frank appearance of proteinopathy. Cross-talk between AD risk immune genes has been recently demonstrated, notably between Cd33 and Trem2 (Griciuc et al., 2019). The TG-Abi3-Gngt2-/- mice also over time showed dysfunctional immune phenotype, with age-dependent reduction of beneficial effect on Aβ and accumulation of ptau. Overall, this would suggest that loss of *Abi3*-*Gngt2* function leads to early induction of immunity that could lead to detrimental outcomes in the longer term (Fig. 11). It is known that phosphorylation is key to maintaining the stability and function of ABI family of proteins (Courtney et al., 2000, Mendoza, 2013). Thus, our data showing that the AD-associated mutation results in hypomorphic function or partial loss of function. Based on our data, it is tempting to suggest that wild type ABI3 could suppress immune gene expression and modulate the cross-talk between different AD immune risk factors that is normally associated AD progression and possibly lead to a beneficial outcome (Fig. 11).

## Materials and Methods

### Mice

All animal studies were approved by the University of Florida IACUC and recombinant DNA use was approved by University of Florida EH&S Office. In this study, we have created several lines of new transgenic *APP* mouse models that are deficient in Abi3. We obtained live *Abi3*(+/-) mice on a BL/6 background from Jax Labs (B6N(Cg)-Abi3tm1.1(KOMP)Vlcg/J; Stock #028180). *Abi3*(+/-) mice were bred to TgCRND8 mice (*APP*swe/ind) (Janus et al., 2000), maintained on BL/6xC3H background, which were then crossed with *Abi3*(-/-) to generate the three different *Abi3* genotypes ( wild type: +/+; heterozygous +/-; knock-out: -/-) with the *APP* transgene. Mice were healthy and without any obvious phenotype, except sudden death of 25% of *APP* transgenic mice (as observed in parental TgCRND8 mice). Mice were maintained on food and water ad libitum on a 12 hour light/dark cycle. All animals were euthanized following IACUC-approved procedures, with AAV expressing animals additionally being perfused intra-cardially post mortem with cold saline. Brains were collected immediately, with left hemisphere flash frozen and right hemisphere drop-fixed in 10% normal buffered formalin. The sex distribution of each cohort is indicated in the figures (open or closed circles) or indicated in figure legends.

### Human tissue specimens

Formalin-fixed, paraffin embedded brain tissue samples of de-identified patients with AD and normal control subjects were provided by the University of Florida Neuromedicine Human Brain and Tissue Bank (UF HBTB) following institutional regulations.

### AAV production and neonatal injection

Recombinant AAV was produced and injected in neonatal mice on day P0 as described earlier (Chakrabarty et al., 2013). Specifically, an AAV construct expressing the 0N/4R human tau construct under the control of CBA promoter (Koller, 2019) was packaged in capsid serotype 1 for these experiments. Control mice were injected with an empty vector (AAV vector backbone CTR0 packaged in serotype 1). All animals were injected with 2µl (1×10^10^ vector genomes) of AAV1 in the cerebral ventricles of each hemisphere as described before (Chakrabarty et al., 2013).

### Biochemical extraction of proteins from brain

Frozen forebrains (without cerebellum or olfactory bulbs) were cryopulverized and split into two groups for *APP* mice, one for Aβ analysis and another for RNA analysis. All ultracentrifugation was done in a Beckman Optima TLX centrifuge with a fixed angle TLA55 rotor. All lysates were prepared to a final concentration of 1ml/150mg of dry tissue. The first aliquot was weighed on dry ice and added to RIPA buffer (50 mM Tris-HCl, 150 mM NaCl, 1% Triton X-100, 0.5% Deoxycholate, 0.1%SDS, 1x protease inhibitor cocktail). Tissues were homogenized using Tissue Master (Omni International Tissue Master 125) for 30 seconds. Tissue homogenate were centrifuged at 43,000 rpm for 1 hour at 4°C. Supernatant was aliquoted and stored at −80 °C as the RIPA lysate. The residual pellet was resuspended in 2% sodium dodecyl sulfate (SDS), sonicated with 3 bursts of 1 minute each (Misonix Q700), and then centrifuged at 43,000 rpm for 1 hour at 15 °C. The supernatant was aliquoted and stored at −80 °C as the SDS lysate. The pellet was resuspended in 70% formic acid (FA), sonicated, centrifuged at 43,000 rpm at 15 °C, and the supernatant stored at −80 °C as the FA lysate.

For AAV-injected brain, the brains were cryopulverized and extracted using tau-specific buffer conditions as described before (Koller, 2019). Briefly, brains were homogenized (Omni International) in appropriate amount of TBS buffer (50mM Tris base, 274 mM NaCl, 5 mM KCl, pH 8) containing protease and phosphatase inhibitors (Pierce Protease & Phosphatase Inhibitor Mini Tablets, ThermoScientific). Following centrifugation at 22,000 rpm for 20 minutes at 4 °C in a Beckman ultracentrifuge, supernatant was stored at −80 °C as S1. The pellet, P1, was resuspended in high salt buffer (10mM Tris base, 0.8M NaCl, and 10% Sucrose) containing 0.05% Triton x-100 (Fisher) and protease and phosphatase inhibitors (Pierce Protease & Phosphatase Inhibitor Mini Tablets, ThermoScientific) and separated at 22,000 rpm for 20 min at 4 °C into S2 and P2. Pellet P2 was resuspended in 1% sarkosyl, incubated at 37 °C for one hour with intermittent gentle shaking and separated at 60,000 rpm for 1hr at 15 °C into S3 and P3. Pellet P3 was resuspended in urea/SDS buffer (4M urea, 2% SDS, and 25mM Tris HCl, pH 7.61) by sonicating (Misonix sonicator) and separated at 50,000 rpm at 15 °C for 30 min. The resulting supernatant S4 was designated as insoluble NFT tau.

### Immunohistochemical and histological analysis

Formalin fixed brains were paraffin embedded, and used for analysis. Slides were deparaffinized using standard procedure (described in (Koller, 2019)) and antigen retrieval performed with steam. Slides were blocked in 2% FBS in 1xPBS for 1 hour at room temperature followed by primary antibody incubation at 4 °C overnight (see **Table EV4**). Appropriate secondary antibody (ImmPress reagents, Vector Labs) was used followed by detection using DAB (Vector Labs) and hematoxylin counterstaining. Slides were mounted in Permount mounting media. ThioS staining was done by incubating deparaffinized and rehydrated slides in 1% ThioS (see **Table EV4**) for 7 minutes at room temperature. Slides were quickly washed in 70% ethanol and then water and mounted with Fluoromount containing DAPI.

### Analysis of Immunochemical and Histological Images

Images of stained slides were captured using Scanscope XT image scanner (Aperio, Vista, CA, USA) and percent area of immunostained slides was quantified with the Positive Pixel Count program (Aperio) that detects DAB staining. 3 section per brain was selected and quantified. Data is presented as % immunoreactivity ± sem and statistical comparisons were conducted using 1-Way ANOVA with a Tukey post hoc test, if necessary. Fluorescent images were captured using BZ-X710 All-in-one Fluorescence Microscope (Keyence Co., Itasca, IL). MC1 stained sections and ThioS stained sections were manually counted for the number of cells or plaques (respectively) positive for the stain.

### Biochemical analysis of tissue lysates

Protein concentrations of RIPA, SDS (in *APP* mice) and S4 fractions (in AAV tau expressing mice) were determined using Bicinchoninic Acid assay (Pierce BCA Protein Assay Kit, ThermoScientific). 20-25 µg of RIPA or SDS lysate or 1 µg of S4 lysate were separated in a 4-20% Tris-glycine gel (Novex, Invitrogen) and transferred to PVDF membranes. Membranes were blocked for 1 hr in 0.5% casein and incubated overnight at 4 °C in primary antibody (see **Table EV4**). Membranes were incubated in appropriate secondary antibody (see **Table EV4**) diluted in 0.5% casein (1:20,000) with 0.005% SDS for 1 hour at room temperature. Membranes were washed in 1X TBS and water, and protein bands were detected using the multiplex Li-Cor Odyssey Infrared Imaging system (Li-Cor Biosciences, Lincoln, NE, USA). Relative band intensity was quantified using ImageJ software (NIH).

For ELISA determination of Aβ levels, Immulon 4HBX plates were coated with 20μg/μl of capture antibody (**Table EV4**) overnight at 4 °C. Plates were washed and blocked in Block Ace (**Table EV4**) at 4 °C overnight. Plates were washed and loaded with SDS lysates or FA lysates (neutralized in Tris buffer) at predetermined dilutions (ranging from 1:300 for 3 month old mice to 1:750 for 6 month old mice). Following overnight incubation, plates were washed and incubated in capture antibody specific for Aβ42 or Aβ40. Following thorough washing, colorimetric assay was developed using TMB solution and detected using Spectramax ELISA reader. The data was analyzed using Softmax program.

### RNA-seq and data analysis

RNA was extracted from frozen, pulverized forebrains using TRIzol reagent (Invitrogen). Extracted RNA was cleaned using RNeasy mini extraction kit with an on-column DNase treatment (Qiagen). RNA quality was quantified with the Qubit RNA HS assay (Invitrogen). RNA quality was checked on an Agilent Bioanalyzer 2100 with the Eukaryote Total RNA Nano chip (**Table EV4**). Total RNA (1 microgram) was used for sequencing library preparation using the Illumina TruSeq RNA library prep with polyA purification (**Table EV4**). Libraries were loaded at equimolar quantities and sequenced on paired-end, 75 bp runs on the Nextseq 500 (Illumina) with a goal of attaining a yield of 30-50 Mb of sequence per sample. RNA extraction and sequencing was performed with an aim to reduce batch effects.

### RNA-seq data analysis

FASTQ alignment, gene counts and differential expression analysis. FASTQ files were aligned against the mouse genome (GRCm38) and GRCm38.94 annotation using STAR v2.6.1a (Dobin, Davis et al., 2013) to generate BAM files. BAM files were used to generate gene counts using Rsamtools (Dani, Wood et al., 2018) and the summarizeOverlaps function with the GenomicAlignments package (Lawrence et al., 2013). Differential gene expression analysis was performed with DESeq2 package using the “DESeq” function with default settings (Love et al., 2014) which fits a generalized linear model for each gene. Subsequent Wald test p-values are adjusted for multiple comparisons using the Benjamini-Hochberg method (adjusted p-value). Pair-wise changes in gene expression levels were examined between groups to identify differentially expressed genes (DEGs). DEGs were defined as an absolute log2Fold Change ≥ 0.5 and an adjusted p-value ≤0.05.

### Cell Type Signatures

Gene lists identifying cell-types within the brain and microglial and astrocytic sub-type were obtained from previously published studies (Chen et al., 2020, Friedman et al., 2018, Hammond et al., 2019, Keren-Shaul et al., 2017, Krasemann et al., 2017, Sala Frigerio et al., 2019, Zhang et al., 2014). Using these gene lists, the geometric mean of the FPKM (fragments per kilobase of exon per million mapped fragments) for genes identified for each cell or cellular sub-type was calculated on a per-animal basis. Group means were calculated, and between group significance values determined by one-way ANOVA. Outliers were removed from the group if their value fell outside of 1.5 times the inter-quartile range (IQR).

### WGCNA

The WGCNA package in R (Langfelder & Horvath, 2008, Langfelder & Horvath, 2012) was used to construct gene correlation networks from the expression data after filtering and removing genes with zero variance. Soft power settings were chosen using the “pickSoftThreshold” function within the WGCNA package. Networks were constructed separately for *APP* transgenic and non-transgenic samples. Adjacency matrices were constructed using expression data and these power setting with the “adjacency” function and a signed hybrid network. Module identification was performed using the “cutreeDynamic” function and a deepSplit setting of 2 with a minimum module size of 30 for all analyses. Modules with similar gene expression profiles were merged using the mergeModules function.

Functional annotation of DEGs, heatmap clusters and WGCNA modules. Gene ontology enrichment analysis was performed with goseq v1.42.0 (Young et al., 2010) to identify gene ontology categories and KEGG pathways that are affected for the given gene lists. For DEGs, up- and down-regulated gene lists were analyzed separately. For WGCNA, gene lists from each module were used as input and GOseq analysis was performed for each module separately. Over-represented p-values were adjusted for multiple comparisons using the Benjamini-Hochberg adjustments for controlling false-discovery rates. An enrichment score was calculated by an observed-over-expected ratio of

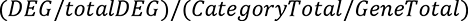

Where DEG represents the total number of DEGs or module genes within the GO or KEGG category, totalDEG represents the total number of DEGs or module genes; CategoryTotal represents the total number of genes within the GO or KEGG category and GeneTotal represents the total number of genes examined. GO terms and KEGG pathways are filtered for p-values adjusted for multiple comparisons (BHadjust) < 0.05, enrichment scores > 1 and total number of genes within the category > 5.

Gene lists to annotate WGCNA modules and identify microglia subtype signatures were identified from previously published studies (Chen et al., 2020, Dejanovic, Huntley et al., 2018, Hammond et al., 2019, Keren-Shaul et al., 2017, Krasemann et al., 2017, Sala Frigerio et al., 2019, Zhang et al., 2014). Gene overlap analysis was conducted with the GeneOverlap package in R (http://shenlab-sinai.github.io/shenlab-sinai/). GeneOverlap uses Fisher’s exact test to calculate the p-value for significance testing as well as calculating the odds ratio. goseq was used for GO and KEGG pathway analysis of genes within each module filtering for those terms with p-values < 0.05, enrichment scores > 1 and total number of genes within the category > 5.

### Correlation analysis from human brains and mouse brains

To assess *GNGT2* and *ABI3* for correlation in human brain, RNAseq expression measures collected from two datasets representing three brain regions (TCX = temporal cortex, CER = cerebellum, DLPFC = dorsolateral prefrontal cortex) were obtained from the AD-knowledge portal (Mayo RNAseq and ROSMAP studies). The AMP-AD consortium previously reprocessed the raw format RNASeq data from these two studies through a consensus alignment, counting and quality control pipeline (RNAseq Harmonization Study). The reprocessed gene counts and associated metadata were downloaded and underwent further quality control, followed by CQN normalization of the raw counts for the 631 ROSMAP, 259 Mayo RNAseq TCX and 246 Mayo RNAseq CER samples that remained. For ROSMAP, four samples were excluded due to missing metadata, one sample removed due to inconsistent sex between provided metadata and expression of Y chromosome genes, and three samples removed based on being outliers (>4SD from mean, PC1 or PC2) in principal components analysis of the reprocessed gene counts (counts per million). For Mayo RNAseq, samples were excluded from the RNASeq datasets based on quality control (QC) outcomes provided in the metadata files. An additional 3 samples were removed due to being gene expression outliers following principal components analysis (>4SD from mean, PC1 or PC2) of the reprocessed gene counts (counts per million). CQN normalized gene counts for *ABI3* and *GNTG2* were extracted for each dataset and plotted in R-3.6.0 using ggplot2. The correlation in the expression between the two genes was assessed using the cor.test function (spearman) in R-3.6.0.

To examine the correlation between Gngt2 and Abi3 in the mouse cortex, bam sequencing alignment files from the TgCRND8 and rTg4510 mouse models were downloaded from the Tau and APP mouse model study at Synapse (10.7303/syn3157182). The gene counting step was performed with the summarizeOverlaps functions in the GenomicAlignments package (Lawrence et al., 2013) and counts were subsequently converted to FPKM with the DESeq2 package (Love et al., 2014) in R 4.1.0. Gngt2 and Abi3 FPKM values were extracted and plotted in R using the ggscatter function and spearman correlation method in the ggpubr package (https://CRAN.R-project.org/package=ggpubr).

### In situ hybridization (RNAscope®, Advanced Cell Diagnostics, Newark, CA) and combined in situ hybridization/immunohistochemistry

For in-situ hybridization, 5µm thick paraffin-embedded tissue sections on slides were rehydrated in xylene and series of ethanol solutions (100%, 90%, and 70%). Slides were incubated with RNAscope® Hydrogen peroxide for 10 min at RT, followed by antigen retrieval in steam for 15 min using RNAscope® 1x target retrieval reagent. After incubation in 10% ethanol for 3 min slides were air dried at 60°C. Subsequently, slides were incubated with RNAscope® Protease plus reagent for 30 min at 40°C in a HybEZ™ oven, followed by 3 washes in distilled water. Slides were then incubated with the following RNAscope® probes (Mouse Abi3 (Cat No. 539161), human ABI3-O1 (Cat No. 549711)) for 2 hours at 40°C in a HybEZ™ oven. Following washes with 1X Wash buffer, slides were incubated with RNAscope®AMP1 solution for 30 min at 40°C followed by series of washes in AMP buffers and incubation in RNAscope® Fast RED-B and RED-A mixture (1:60 ratio) for 10 min at RT. For immunohistochemistry double labeling, sections were incubated in Impress® horse serum (2.5%, Vector Laboratories) solution for 20 min, followed by a 5 min incubation in 2% FBS/0.1 M Tris, pH 7.6. Primary antibodies were diluted in 2% FBS/0.1 M Tris, pH 7.6 at the following dilutions: Iba1 (Wako), 1:500. Sections were incubated with primary antibody over night at 4°C, washed two times in 0.1 M Tris, pH 7.6 for 5 min each and incubated in ImmPress® anti-rabbit IgG plus reagent (Vector Laboratories) for 30 min at RT. After two washes in 0.1 M Tris, pH 7.6 for 5 min each, immunocomplexes were visualized using the chromogen 3,3′-diaminobenzidine (DAB kit; KPL, Gaithersburg, MD). Tissue sections were counterstained with hematoxylin (Sigma Aldrich, St. Louis, MO), air dried at 60°C for 15 min and cover slipped using EcoMount™ mounting medium (Biocare Medical).

### In vitro overexpression and phosphorylation analysis of ABI3

Plasmid encoding for ABI3 (NM_016428) was purchased from OriGene (Catalog# RC202853). Site directed mutagenesis in ABI3 was performed and resulting clones were Sanger sequenced to confirm the presence of mutation. HEK293T cells were grown in DMEM media supplemented with 10% fetal bovine serum (Gibco) and 1% penicillin/streptomycin (Life Technologies) and transiently transfected by CaCl_2_ method. 48 hours later, the transfected cells were lysed in RIPA buffer (Fisher) supplemented with complete EDTA-free Protease Inhibitor Cocktail and phosphatase inhibitor tablets (Sigma-Aldrich). Protein extracts were loaded on Bis-Tris precast gels (BioRad) were transferred on PVDF membrane for Western blotting with the Li-Cor system. ABI3 was detected with the following antibodies according to the experiment: FLAG-M2 (Sigma-Aldrich), N terminal Anti-ABI3 (Sigma-Aldrich) or anti-ABI3 209F (manufactured by Pacific Immunology Corp using the peptide sequence 195-PVVPDGRLSAASSAF-209). Lambda Protein Phosphatase (NEB) assay was performed as described by the supplier. Briefly, ABI3 HEK transfected cells were lysed in PBS 0.1% NP-40 in the presence of complete protease inhibitor. Cytoplasmic proteins fractions were used for the reaction and analyzed by Western Blotting.

### Statistical Analysis

Detailed description of statistics is provided in each figure legend and in the methods section. Statistical analysis was done using GraphPad Prism.

## Acknowledgments

We thank the State of Florida Ed & Ethel Moore Alzheimer’s Disease Research Program (8AZ16), NIH/NIA (U01 AG046139), University of Florida Neuromedicine Human Brain and Tissue Bank (funded in part by NIA P30AG066506) and McKnight Brain Institute for support. We posthumously thank Dr. Peter Davies for anti tau antibodies.

## Author Contributions

PC conceived the study and wrote the manuscript; KRI performed immunohistochemistry, ELISA and western blots and assembled final figures; KNM extracted RNA, performed RNAseq and analyzed data; JL, TEG and SP performed in situ hybridization; MA, XW, ZQ, CF, TEG and NET performed informatics analysis on human AMP-AD data; CL performed in vitro experiments; LZ, SS and QV performed immunostaining and data analysis; EGDLC euthanized animals; PC maintained mouse colonies and performed neonatal injections; TEG and DR provided recombinant AAV. All authors read and reviewed the manuscript.

## Data and Code Availability

The full datasets and R codes supporting the current study are available from the corresponding author on request. On publication, these data will be deposited in AD Knowledge Portal (https://adknowledgeportal.org). The AD Knowledge Portal is a platform for accessing data, analyses, and tools generated by the Accelerating Medicines Partnership (AMP-AD) Target Discovery Program and other National Institute on Aging (NIA)-supported programs to enable open-science practices and accelerate translational learning. The data, analyses and tools are shared early in the research cycle without a publication embargo on secondary use. Data is available for general research use according to the following requirements for data access and data attribution (https://adknowledgeportal.org/DataAccess/Instructions).

## Expanded View (Supplementary) Figure Legends

**Fig. EV1. Localization and levels of ABI3 RNA in human and mouse**.

In situ hybridization was done to detect Abi3 RNA on human (a-f) and mouse (g-x) paraffin embedded brain sections. Human or mouse ABI3 specific RNAscope probes were used for in situ hybridization detected by Fast Red (red color) followed by immunohistochemistry for Iba-1 antibody detected by DAB (brown color). Three representative forebrain images in the vicinity or away from amyloid plaques are shown. Asterisks mark Aβ deposits; arrowheads indicate Iba-1 (microglia) associated in situ signal and arrows indicate in situ signal in non-microglia cells. n=3 (human AD cases, 6 month old TG-*Abi3-Gngt2*^-/-^ mice and 6 month old TG-*Abi3-Gngt2*^+/+^ mice mice) and n=1 (20 month old TgCRND8 mice, collected independent of this study). Representative of two independent experimental replicates. TG= transgenic CRND8. Also see Fig. 1.

**Fig. EV2. Genomic organization at the *Abi3* locus of Mus musculus.**

a. RNAseq based confirmation of Abi3, Gngt2 and APP levels from three *Abi3* genotypes (WT +/+, heterozygous +/-, homozygous -/-) in *APP* transgenic mice or NonTG (*APP* nTG) littermates. b. The *Abi3* knock-out mice (Abi3tm1.1(KOMP)Vlcg) was generated by cre-mediated excision of the parental Abi3tm1(KOMP)Vlcg allele resulting in the removal of the neomycin selection cassette, leaving behind the inserted lacZ reporter sequence. The sequence that was excised out on chromosome 11 was located between 95842143 and 95832627.

**Fig. EV3. Hub genes characteristic of WGCNA modules identified in 3 month old *Abi3-Gngt2*^-/-^ mice.**

a. Volcano plot and table of altered genes representing DEG in 3 month old *Abi3-Gngt2*^+/-^ vs *Abi3-Gngt2*^+/+^ mice. FC, fold change; padj, adjusted p value. b-g. Module membership is plotted against gene connectivity (kWithin) for genes identified within modules significantly correlated with *Abi3-Gngt2* genotype and other neuropathology traits in TG-*Abi3-Gngt2*^-/-^ mice. The top 10 hub genes (as ranked by kWithin values) identified in each co-expression WGCNA modules of 3 month old *Abi3-Gngt2*^-/-^ mice relative to *Abi3-Gngt2*^+/+^ mice. The module members of antiquewhite2, coral2 and mediumpurple2 modules are shown (b, d, f), with top hub genes tabulated (c, e, g). Module statistics are denoted by: kWithin, extent of gene connectivity within the module; MM, module membership value (of gene to module); GS, gene significance value to specific experimental trait. H. The overlap of genes within WGCNA modules with genes previously identified in microglial and astrocytic sub-types is expressed as odds ratio value. The different cell signatures are: neurotoxic A1 and neurotrophic A2 astrocyte (Liddelow et al., 2017); PIG network (Chen et al., 2020); DAM (Keren-Shaul et al., 2017); MGnD and homeostatic microglia (Krasemann et al., 2017). Higher odds ratio denotes higher correlation. All p values are adjusted for multiple testing. Grey boxes indicate non-significant odds ratio values (p-value < 0.5). N=4 mice (2 male, 2 female) per cohort.

**Fig. EV4. Neuropathological attributes of TG-*Abi3-Gngt2*^-/-^ mice.**

a-b. Anti CT20 immunoblot indicating full length APP and C terminal fragments (CTF) in *APP* transgenic mice with intact (+/+), partial loss (+/-) or complete loss (-/-) of *Abi3-Gngt2* (a). Lane marked with asterisk denotes a non transgenic mice for *APP* (a). APP levels normalized to actin is depicted (b). Ubiquitin decorating Aβ plaques in 3 month (c) and 6 month old (d) *APP* TG mice with intact (+/+), partial loss (+/-) or complete loss (-/-) of *Abi3*-*Gngt2*. Scale Bar, 2mm (whole brain), 50 µm (cortex and hippocampus). e-m. Representative brain images stained with 33.1.1 antibody and corresponding quantitation (e-f), GFAP antibody and corresponding quantitation (g-h), Iba-1 antibody and corresponding quantitation (i-j), cd11b and corresponding quantitation (k-l) and ubiquitin decorated Aβ plaques (m) in 9 month old *APP* TG mice with intact (+/+), or partial loss (+/-) of *Abi3*-*Gngt2*. Scale Bar, 2 mm (whole brain), 50 µm (cortex and hippocampus). N=7-8 mice/genotype. Data represents mean±sem.

**Fig. EV5. Synaptic protein levels in TG-*Abi3-Gngt2*^-/-^ mice.**

Immunoblotting depicting levels of synaptic proteins in 3 month old *APP* TG mice with intact (+/+), partial loss (+/-) or complete loss (-/-) of *Abi3-Gngt2* (a, c, e, g, i, k, m). Quantitation of the synaptic proteins normalized to actin or GAPDH is depicted (b, d, f, h, j, l, n). Molecular weight markers in kDa are indicated on each panel. Data represents mean±sem. 1 way Anova; **p<0.01; *p<0.05.

**Fig. EV6. Hub genes characteristic of WGCNA modules identified in 3 month old TG-*Abi3-Gngt2*^-/-^ mice.**

a. No significant genes were identified in 3 month old TG-*Abi3-Gngt2*^+/-^ mice relative to TG-*Abi3-Gngt2*^-/-^ mice. FC, fold change; padj, adjusted p value. b-g. Module membership is plotted against gene connectivity (kWithin) for genes identified within modules significantly correlated with *Abi3*-*Gngt2* genotype and other neuropathology traits in TG-*Abi3-Gngt2*^-/-^ mice. The top 10 hub genes (as ranked by kWithin values) identified in each co-expression WGCNA modules of 3 month old TG-*Abi3-Gngt2*^-/-^ mice relative to TG-*Abi3-Gngt2*^+/+^ mice. The module members of honeydew1, plum3 and sienna3 modules are shown (b, d, f), with top hub genes tabulated (c, e, g). Module statistics are denoted by: kWithin, extent of gene connectivity within the module; MM, module membership value (of gene to module); GS, gene significance value to specific experimental trait. h. The overlap of genes within WGCNA modules with genes previously identified in microglial and astrocytic sub-types signatures is expressed as odds ratio value. The different cell signatures are: neurotoxic A1 and neurotrophic A2 astrocyte (Liddelow et al., 2017); Plaque-induced gene (PIG) network (Bagnat, Keranen et al., 2000); Disease-associated microglia (DAM) (Keren-Shaul et al., 2017); microglial neurodegenerative phenotype (MGnD) and homeostatic microglia (Krasemann et al., 2017). All p values are adjusted for multiple testing (padj). Higher odds ratio denotes higher correlation. Grey boxes indicate non-significant odds ratio values (p-value < 0.5). 4 mice (2 male, 2 female) per cohort.

**Fig. EV7. Somatic transgenesis modeling of tauopathy in *Abi3-Gngt2*^-/-^ mice.**

a-c. Mice were injected with control vector (Control) or AAV expressing a double mutant (P301L/S320F) human tau on neonatal day P0 and analyzed at 3 months or 6 months. a. Representative images of ThioS-stained cortex of *Abi3-Gngt2*^+/+^ or *Abi3-Gngt2*^-/-^ mice injected with AAV-P301L/S320F tau or control vector shown. Three individual mice from each cohort are shown. b-c. Guanidine-insoluble brain lysates of AAV-P301L/S320F tau expressing *Abi3-Gngt2*^+/+^ or *Abi3-Gngt2*^-/-^ mice were separated on an immunoblot and probed with CP27 antibody (b) or CP13 antibody (c) to show presence of insoluble NFT tau. N=3 mice/condition. d-s. Mice were injected with control vector (Control) or AAV expressing wild type human tau on neonatal day P0 and analyzed at 3 months or 6 months. Representative brain images of total human tau (detected with CP27 antibody, d, l), ptau (detected with CP13 antibody, f, n), astrocytes (detected with GFAP antibody, h, p) and microglia (detected with Iba-1 antibody, j, r) from cortex and hippocampus of *Abi3-Gngt2*^+/+^ or *Abi3-Gngt2*^-/-^ mice are shown. Quantitative analysis of antibody-stained brain sections (e, g, i, k, m, o, q, s) shown below corresponding stained brain images (d, f, h, j, l, n, p, r). Quantitative data in the graphs depicting the control vector cohort (*Abi3-Gngt2*^+/+^ or *Abi3-Gngt2*^-/-^ genotypes) is shared with corresponding data in Fig. 8 and Fig. 9 as these experiments were done simultaneously. For representative images of the control cohort, please refer to Fig. 8 and Fig. 9. Scale bar, 75 µm. n=4-6 mice/group. Data represents mean±sem. 1 way Anova; ****p<0.0001; ***p<0.001; **p<0.01; *p<0.05.

## Expanded View Table Legends

**Table EV1.** Normalized RNA levels (FKPM values) of Abi3 and Gngt2 from TgCRND8 mice at various ages

**Table EV2.** Normalized RNA levels (FKPM values) of Abi3 and Gngt2 from rTg4510 mice at various ages

**Table EV3**. Analysis of Abi3 and Gngt2 expression changes in mouse studies (transcriptomic data originally reported by Wan et al. [32] and Al Ouran et al [33] and available at http://mmad.nrihub.org)

**Table EV4.** Description of resources and reagents used in this study.

## Abbreviations

+/+: wild type for *Abi3*-*Gngt2*
+/-: haploinsufficient for *Abi3*-*Gngt2*
-/-: complete deficiency of *Abi3*-*Gngt2*
AAV: adeno-associated virus
Aβ: amyloid β
ABI3: Abelson Interactor Protein 3
AD: Alzheimer’s disease
Adamts6: ADAM Metallopeptidase With Thrombospondin Type 1 Motif 6
Adgrf3: Adhesion G Protein-Coupled Receptor F3
Aklr1c13: Aldo-keto reductase family 1 member C13
AMP-AD: Accelerating Medicines Partnership
ARM: Activated response microglia
CER: cerebellum
Chil1: Chitinase-like 1
Clec7a: C-type lectin domain family 7 member A
Cxcr4: C-X-C Motif Chemokine Receptor 4)
DAM: damage associated microglia
DEG: differentially expressed genes
DLPFC: dorsolateral prefrontal cortex
FC: fold change
Fndc9: Fibronectin Type III Domain Containing 9
FPKM: fragments per kilobase of exon per million mapped fragments
GO: Gene Ontology
GFAP: glial fibrillary acidic protein
Gngt2: Guanine Nucleotide Binding Protein Gamma Transducing Activity Polypeptide 2
GS: gene significance value to specific experimental trait
Iba-1: ionized calcium-binding adapter molecule 1
Irf8: Interferon Regulatory Factor 8
Ly86: Lymphocyte Antigen 86
kWithin: extent of gene connectivity within the module
MAF: minor allele frequency
MM: module membership value
MGnD: microglial neurodegenerative phenotype
Mpeg1: Macrophage Expressed 1
MAPT: microtubule associated protein tau
Nramp1: Natural resistance-associated macrophage protein 1
NESH: new molecule including SH3
NFT: neurofibrillary tangles
NonTG: non-transgenic for *APP*swe/ind (CRND8 line)
ns: non-significant
OR: Odds ratio
Padj: p values adjusted for multiple comparisons
Pcdha5: Protocadherin Gamma Subfamily A 5
PIG: plaque induced genes
QC: quality control
RNA: sequencing RNAseq
S100a9: S100 Calcium Binding Protein A9
SDS: sodium dodecyl sulfate
SH3: Src homology 3
Sval1: seminal vesicle antigen-like 1
TCX: temporal cortex
ThioS: thioflavin S
TG: Transgenic for swe/ind (CRND8)
WGCNA: weighted gene co-expression network analysis
WT: wild type

